# Identification and Characterization of a Small Molecule Ligand for the Huntingtin-HAP40 Complex

**DOI:** 10.1101/2025.11.21.689733

**Authors:** Rocher Leung, Esther Wolf, Justin C. Deme, Swati Balakrishnan, Conrad V. Simoben, Shulu Feng, Jonathan D. Macdonald, Renu Chandrasekaran, Jianxian Sun, Xiaoyun Wang, Albina Bolotokova, Yefen Zou, Ram Kannan, Suzanne Ackloo, Derek Wilson, Susan M. Lea, Aled M. Edwards, Cheryl H. Arrowsmith, Hui Peng, Levon Halabelian, Matthieu Schapira, Rachel J. Harding

## Abstract

Huntington’s disease (HD) is caused by a CAG repeat expansion mutation, giving rise to a polyglutamine expansion in the huntingtin (HTT). However, the explicit molecular functions of HTT and opportunities for direct pharmacological modulation remain incompletely understood. Here, we report the discovery of a small molecule ligand for the full-length HTT protein in complex with its partner, HAP40. Using affinity selection mass spectrometry (AS-MS), we identified a stereoselective binder, whose binding was characterized by surface plasmon resonance, hydrogen-deuterium exchange mass spectrometry, and cryo-electron microscopy at 2.3 Å resolution. The ligand binds HTT-HAP40 *in vitro* with single-digit micromolar affinity and one-to-one stoichiometry at a druggable interface previously predicted computationally. *In silico* studies predicted and experimental analyses confirmed the (R)-enantiomer as the eutomer and initial structure activity relationship was established experimentally. This work details a structurally-validated chemical scaffold and highlights a ligandable pocket which could enable development of chemical probes for probing HTT biology, as well as therapeutics such as degraders and imaging agents for HD.

## Introduction

Huntington’s disease (HD) is an inherited neurodegenerative disorder caused by a CAG trinucleotide repeat expansion in the *Huntingtin* (*HTT*) gene, leading to the expression of a mutant huntingtin protein with an expanded polyglutamine (polyQ) tract.^1^ HD is a monogenic and fully penetrant disorder, however, the underlying molecular mechanisms of disease progression remain incompletely understood.^2^ Hallmark features of HD pathology, including protein aggregation,^3,4^ transcriptional dysregulation,^5,6^ and somatic expansion,^7^ track with disease onset and progression, and delineating the causal roles and relative contributions of these phenomena to the cascade of progressive symptoms experienced by patients remains a key focus of the HD research field.

To date, most clinical strategies for HD have focused on reducing HTT protein levels using splice modulators, antisense oligonucleotides, or viral-delivered gene therapies.^8^ However, no disease-modifying therapy has yet been approved.^9^ Alternative therapeutic strategies that target the HTT protein directly, rather than suppressing its production might overcome some barriers faced by these approaches that predominantly target HTT mRNA.

High-quality small molecule ligands are powerful tools for probing protein function and validating therapeutic hypotheses.^10,11^ HTT has long been considered a challenging target for medicinal chemistry campaigns due to its large size (3144 amino acids), absence of enzymatic activity, and conformational flexibility.^12,13^ Prior efforts to develop HTT-targeting molecules have largely focused on aggregated exon 1 fragments using flat, multicyclic compounds that often lack specificity and fail to bind full-length, soluble HTT.^14,15^ Similarly, attempts to image or degrade mHTT aggregates with radioligands or bifunctional degraders respectively have been confounded by poor selectivity, non-specific fibril binding, or an absence of ideal pharmacological hallmarks such as hook effects in proteolysis-targeting chimera (PROTAC)-style molecules.^14–17^

Here, we report the discovery of a small molecule ligand that directly and specifically binds full-length HTT in complex with its stabilizing partner HAP40.^18^ The HTT-HAP40 complex represents a dominant physiological form of soluble HTT in human cells, and also is a tractable protein with optimal biophysical properties for a ligand discovery campaign.^18,19^ Building on our prior identification of macrocyclic peptides that bind HTT with high affinity and selectivity,^20^ we combined affinity selection mass spectrometry (AS-MS), surface plasmon resonance (SPR), hydrogen-deuterium exchange mass spectrometry (HDX-MS), and cryo-electron microscopy (cryo-EM) to discover, validate, and structurally characterize a series of small molecule HTT ligands. One binder interacted with full-length HTT with one-to-one stoichiometry and single-digit micromolar affinity in a druggable pocket we previously predicted at the interface of the HTT-HAP40 complex.^18^ We defined a preliminary structure-activity relationship (SAR) for this chemical scaffold and found chemical analogs with improved binding affinity and comparable aqueous solubility. Thus, this ligand is tractable for structure-guided optimization and development into tools (e.g., tracers) and therapeutics (e.g., degraders).

## Results and Discussion

### Hit-finding using affinity selection mass spectrometry (AS-MS)

Previously, our lab purified the HTT-HAP40 complex, yielding a highly pure, monodisperse and structurally validated sample.^18,21,22^ In this study, HTT constructs with a polyglutamine tract of either 23 or 54 residues (Q23 and Q54, respectively) were studied. We screened HTTQ54-HAP40 (referred to as HTT-HAP40) against a 15K library of drug-like compounds spanning over 7000 scaffolds using AS-MS (**Fig 1a**).^23^ Several hits, such as **AS-MS Hit 1** (**1**) and **AS-MS Hit 2** (**2**) (**Fig 1b**), were enriched for this complex compared to at least seven unrelated proteins screened in parallel. These hits did not bind 101 other proteins that were screened retrospectively under the same conditions, demonstrating their selective binding to HTT-HAP40. For **AS-MS Hit 1**, we used chiral chromatography to compare the ratio of the two enantiomers in both the chemical library stock and in the eluate from HTT-HAP40. The bound eluate sample was enriched for one enantiomer, indicating enantiomeric selectivity in binding of **AS-MS Hit 1** to HTT-HAP40 (**Fig 1c**).

**Figure 1.**
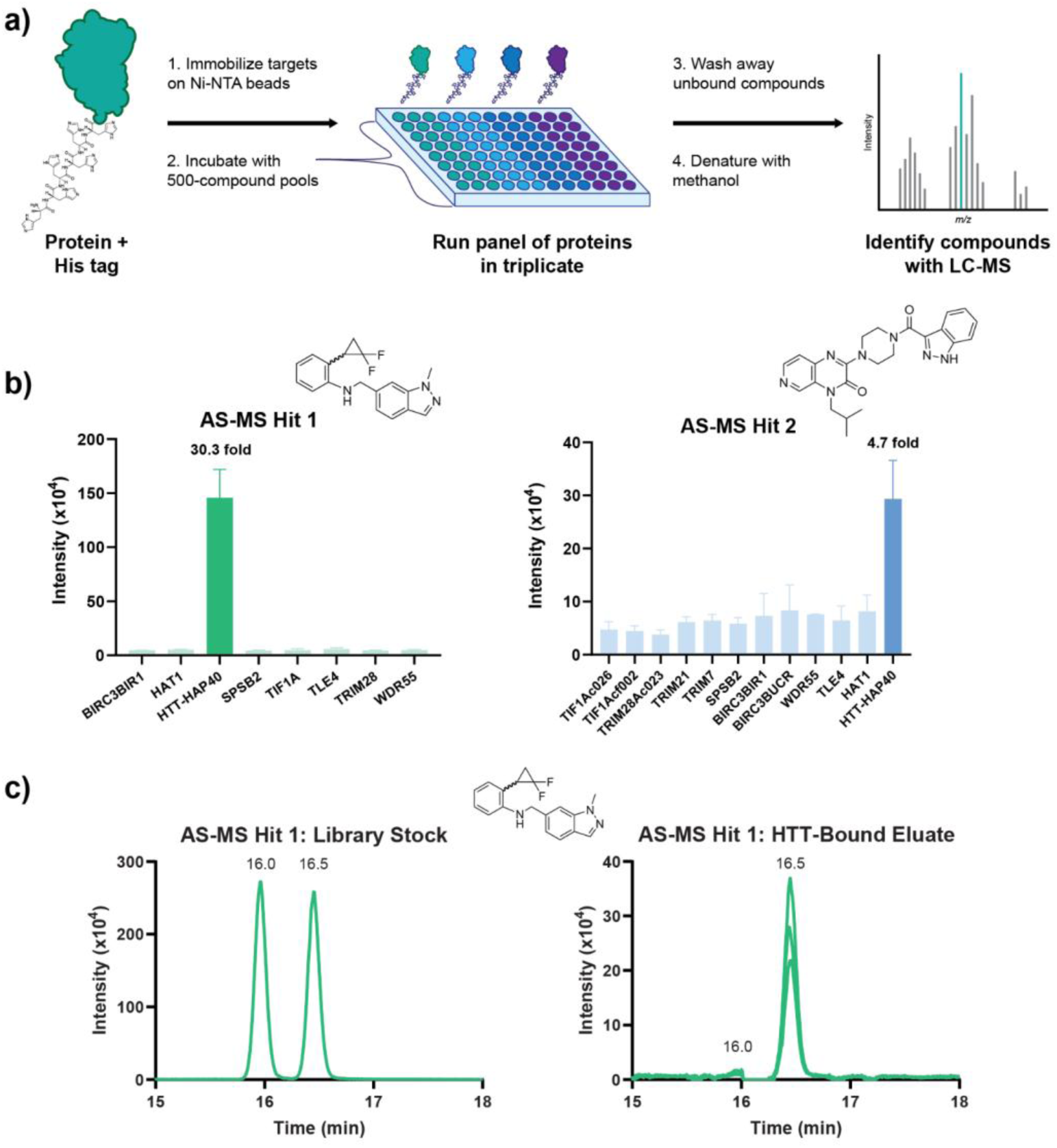
AS-MS screening of small molecules against HTT-HAP40. **a)** Schematic representation of the AS-MS screening workflow. **b)** Examples of hits enriched for HTT-HAP40. Fold change was calculated as the average signal intensity for HTT-HAP40 divided by the average signal intensity (N=3) across all other tested proteins. Error bars represent standard deviation. **c)** Chiral separation was performed on **AS-MS Hit 1** to compare the ratio of enantiomers present in the chemical library with that of the HTT-bound eluates (N=3) taken from the AS-MS screen.

### Orthogonal confirmation and characterization of HTT-HAP40 binding of AS-MS hits

To confirm binding of the putative AS-MS hits, top-ranking compounds were purchased as pure solids for testing in an orthogonal assay. SPR was performed using biotinylated HTT-HAP40 protein, immobilized on a streptavidin-coated sensor chip, and each compound was titrated to assess binding. Two compounds demonstrated dose-dependent binding with affinities in the low micromolar range (**Fig 2ab**). One of these hits, **AS-MS Hit 1** (**1**), had an average apparent K_D_ of 5 ± 2 μM and an average %R_max_ of 110% ± 12% (N>10) (**Fig 2a, Table S1**). The %R_max_ was calculated by normalizing the experimental maximum response to the theoretical maximum response of stoichiometric (1:1) binding for each immobilized ligand and analyte.^24,25^ The %R_max_ of **AS-MS Hit 1** was consistent with a stoichiometric binding profile and suggested specific binding to HTT-HAP40.

**Figure 2.**
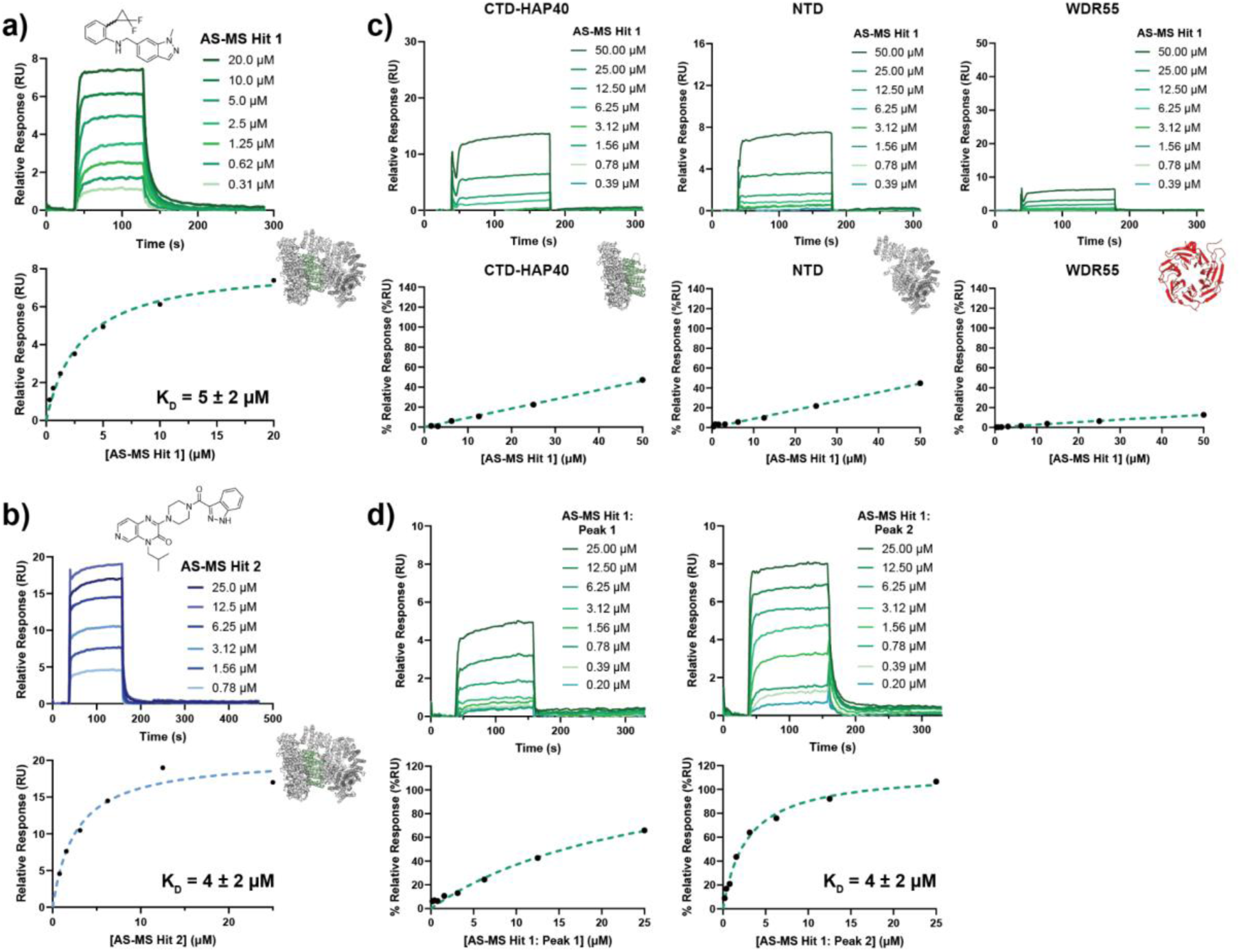
*AS-MS hit validation and characterization of binding to HTT-HAP40 by SPR*. Validation of dose-response for **a) AS-MS Hit 1** and **b) AS-MS Hit 2** through steady state response (black circles) and a 1:1 binding model fit (dashed green line). Raw sensorgrams and affinity plots are representative of at least three replicates and K_D_ is reported as mean ± standard deviation (N≥3). **c)** Characterization of binding for **AS-MS Hit 1** against various HTT subdomains and a negative control protein, WDR55, through the steady state response (black dots) 1:1 binding model (dashed line). Raw sensorgrams and affinity plots are representative of at least three replicates performed with the HTT subdomains and WDR55 (N≥3). K_D_ is considered undetermined if the steady state response plot does not fully reach saturation and the K_D_ is greater than half the maximum analyte concentration tested. **d)** Enantiomeric selectivity of binding for Peak 2 of **AS-MS Hit 1** chiral separation is observed through steady state response (black circles) and a 1:1 binding model fit (dashed green line) with representative sensorgrams, affinity plots, and K_D_ values (N≥8).

To further interrogate the binding profile of **AS-MS Hit 1**, we used a panel of HTT variants, in addition to an unrelated protein, WDR55, to investigate its binding specificity. Our HTT panel included structure-rationalised subdomain constructs which bisect the HTT-HAP40 complex into the N-terminal domain (NTD, aa. T97-M2069) and the C-terminal domain (CTD, aa. V2095-V3138) bound to HAP40.^22^ **AS-MS Hit 1** exhibited dose-dependent binding to the HTT-HAP40 complex, but not the subdomain proteins or WDR55, suggesting specific binding to an interface only present in the full-length protein complex (N=3) (**Fig 2c**).

SPR analysis of **AS-MS Hit 1** in the context of HTT-HAP40 bearing different polyglutamine tracts revealed that binding affinity was indistinguishable between wild-type (Q23) HTT-HAP40 and its expanded form (Q54) (**Fig S1**). The lack of polyglutamine length-dependent effects on ligand binding suggest that ligand recognition is driven by conserved structural features of the HTT-HAP40 complex rather than polyglutamine expansion.

The second hit, referred to as **AS-MS Hit 2** (**2**), had an average apparent K_D_ of 4 ± 2 μM, but showed super-stoichiometric binding by its average %R_max_ of 230% ± 75% (N=3) (**Fig 2b, Table S1**). This suggested non-specific binding or potential aggregation of the compound. **AS-MS Hit 2** also exhibited a linear dose-response with other HTT subdomains and an unrelated protein (WDR55), suggesting the presence of non-specific interactions (**Fig S2**). Its steady-state K_D_ and %R_max_ values across all the tested HTT subdomains and WDR55 could not be determined as its dose-response curves did not reach saturation (N=3) (**Table S1**). Due to the superstoichiometric interactions exhibited across all tested proteins, characterization of **AS-MS Hit 2** was discontinued.

**AS-MS Hit 1** contains a chiral centre and was identified and provisionally validated in its racemic form in both the AS-MS and SPR assays. To further characterise its binding, we separated the racemic mixture into enantiomerically pure compounds, referred to as Peak 1 and Peak 2. SPR analysis of these stereopure compounds and the racemate with HTT-HAP40 revealed binding preferences for Peak 2 which had an average apparent K_D_ of 4 ± 2 μM and an average %R_max_ of 100% ± 9% (N=14) (**Fig 2d, Table S1**) while Peak 1 exhibited a linear dose-response.

### Mapping the binding site of AS-MS Hit 1

SPR analysis of **AS-MS Hit 1** with our suite of HTT subdomains revealed that binding necessitated the full-length HTT-HAP40 complex (∼390 kDa). To identify the binding interface of **AS-MS Hit 1** with HTT-HAP40 we used bottom-up differential HDX-MS (ΔHDX-MS).^26^

The cumulative ΔHDX over 0.25, 1 and 10 minutes was measured to assess the impact of small molecule binding on HTT-HAP40 conformational dynamics at the peptide-level (**Fig 3, S3-S5**). Consistent with our biophysical data and the presumed specificity for the HTT-HAP40 complex, the ΔHDX-MS was localized to a single binding site at the interface of the N-terminal domain of HTT and HAP40, with decreased deuterium uptake spanning residues 1016-1028 and 1073-1085 in HTT and 82-87, 89-93, 122-127, and 133-143 in HAP40. Kinetic analysis revealed that HTT peptide 1076-1085 (LSSAWFPLDL) underwent the highest magnitude and most persistent reduction in deuterium uptake over the time course, suggesting key protein-ligand interactions were being formed (**Fig S5**). On the other hand, HAP40 peptides in the binding site only underwent divergence at the longest timepoint (10 min), suggesting that **AS-MS Hit 1** reduced their conformational freedom. Interestingly, this pocket was identified by our team in a previous study where *in silico* analyses predicted it to be a druggable binding site.^18^

**Figure 3.**
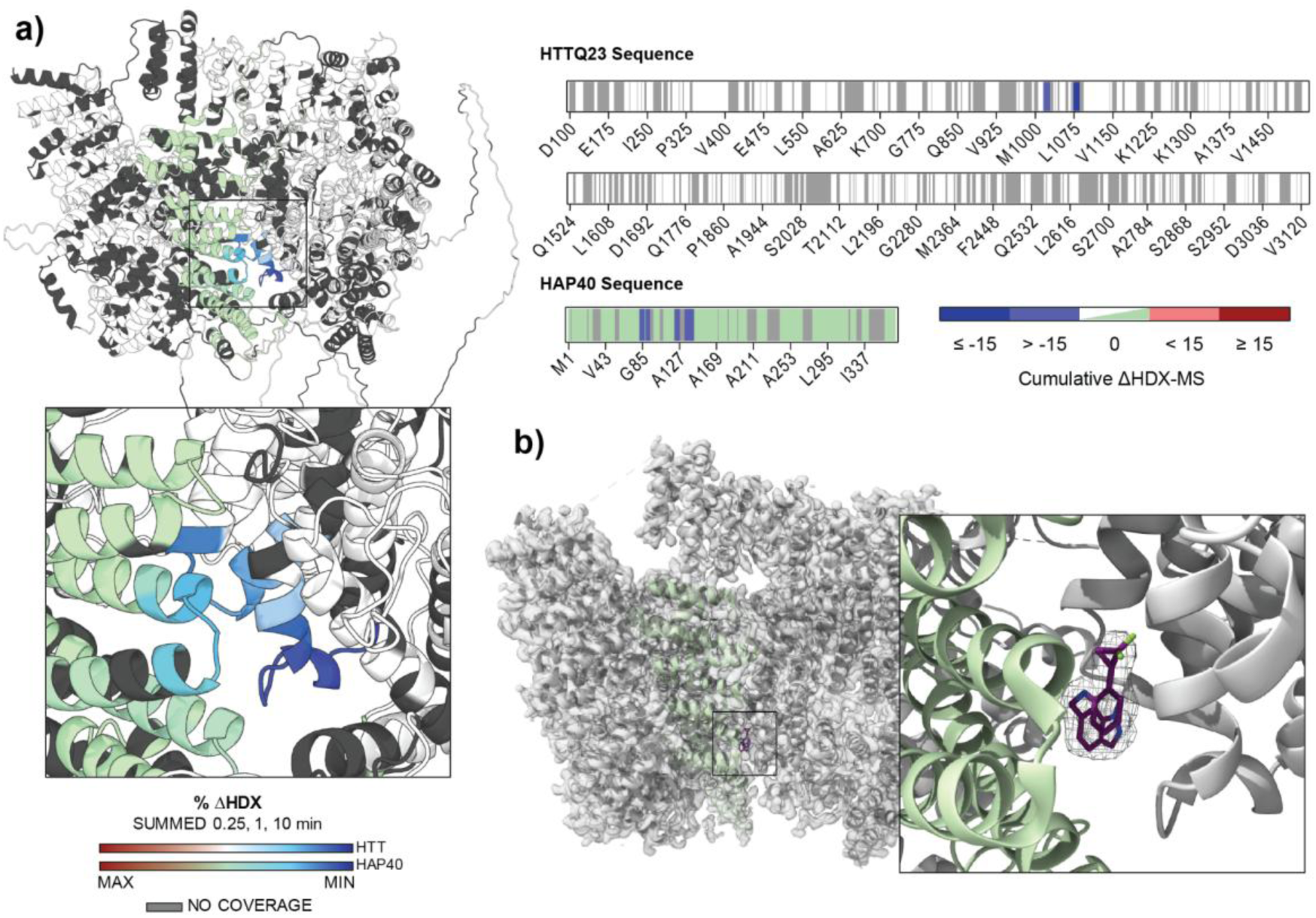
HDX-MS identifies the binding interface of AS-MS Hit 1. **a)** Cumulative per-residue ΔHDX-MS heatmap of HTTQ23-HAP40 and **AS-MS Hit 1**. Statistically significant decreases (blue) and increases (red) are indicated on HTT (white) and HAP40 (green) structures and linearized sequences. Areas lacking peptide sequence coverage are indicated by grey. **b)** Cryo-electron microscopy map reveals binding pocket of **AS-MS Hit 1** (purple) in agreement with HDX-MS data.

To more precisely characterise the binding pose of the ligand in this pocket, we determined the structure of HTT-HAP40 in complex with **AS-MS Hit 1** by cryo-EM. The map was refined to a resolution of 2.3 Å (PDB 9YR6) and the HDX results guided us to the ligand density for which strong map features were observed (**Fig 3ab, S6, Table S2**).

### Determination of the active enantiomer of AS-MS Hit 1

The cryo-EM sample was prepared by saturating the HTT-HAP40 complex with a molar excess of the racemic form of **AS-MS Hit 1**, and the map appeared to accommodate modelling of both enantiomers (**Fig 4a, S6, Table S2**). Based on density fit analysis through Coot,^27^ the (R)-enantiomer fit the map features in the binding pocket more closely, with a density correlation score of 0.64, compared with the much lower score of 0.48 for the (S)-enantiomer.

**Figure 4.**
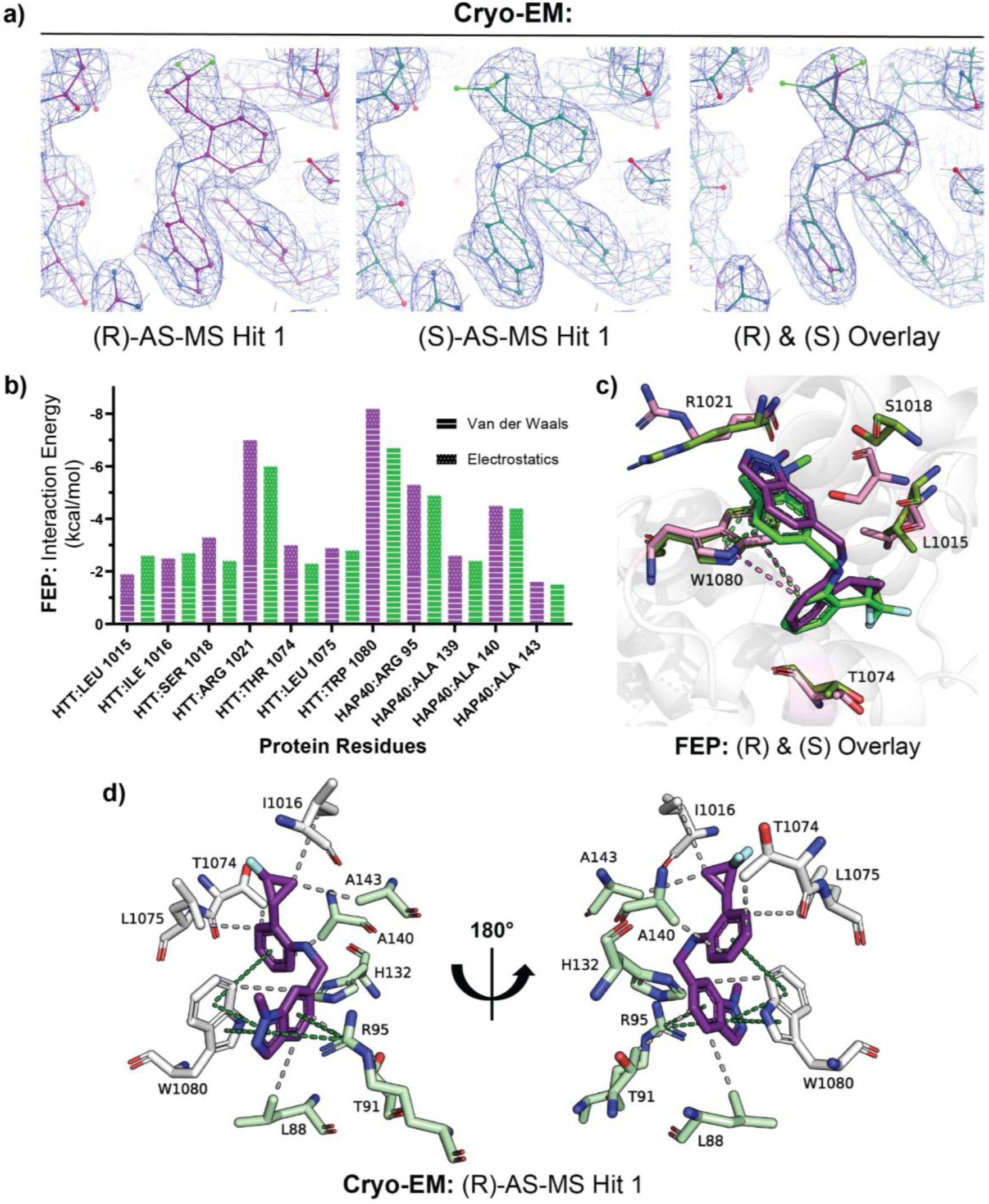
Binding interactions of AS-MS Hit 1 with HTT-HAP40 complex by cryo-EM and FEP. **a)** The density of the cryo-EM map is seen to be strong for the ligand and the binding pocket, with the (R)-enantiomer (purple) having a greater density correlation score than the (S)-enantiomer (green). **b)** Receptor’s per-residue interaction energy. Purple bars represent interactions of the (R)-enantiomer, green bars represent that of the (S)-enantiomer. **c)** Representative binding poses from FEP+ calculation of the enantiomers of **AS-MS Hit 1**. Residues that contribute to the potency difference are highlighted: S1018, R1021, T1074, W1080 and L1015. Corresponding π-interactions and binding site residues are shown in pink for the (R)-enantiomer and green for the (S)-enantiomer. **d)** Cryo-EM binding pose of the (R)-enantiomer of **AS-MS Hit 1** (purple) shows that it binds using a combination of interactions shown in dashed lines: hydrophobic interactions (grey) and π-interactions (green). Residues from HTT are shown in white and those from HAP40 are shown in green.

To characterize the stereochemistry effects and identify potential vectors for further potency improvements, we performed free energy calculations (FEP+) on the two stereoisomers.^28,29^ The binding free energy difference was 1.5 ± 0.1 kcal/mol, with the (R)-enantiomer as the eutomer and the (S)-enantiomer as the distomer. The five residues contributing most to the computed potency differences were S1018, R1021, T1074, W1080 and L1015, all in HTT (**Fig 4bc**). W1080 showed the largest difference of 1.5 kcal/mol and was one of two residues which interacted with the ligand >50% of the simulation time (**Fig 4b**).

W1080 was predicted to form π-π interactions with the phenyl (edge-to-face) and the indazole group (face-to-face) in both stereoisomers. Simulations showed the edge-to-face interaction between the phenyl group and W1080 to be more affected by chiral inversion. The plane of the phenyl group in the (R)-enantiomer was perpendicular to W1080. Conversely, for the (S)-enantiomer, the phenyl group was rotated, decreasing this perpendicular alignment to W1080 and weakening this edge-to-face interaction. This difference was caused by the orientation preference of the difluoro cyclopropane group (**Fig 4c**).

For both stereoisomers, the two fluorine atoms occupied a similar hydrophobic vicinity of the binding site. For the (R)-enantiomer, this placed the cyclopropane group opposite the phenyl plane relative to W1080, favouring π-π interaction with W1080 (**Fig 4d**; PDB 9YR6). In contrast, for the (S)-enantiomer, the cyclopropane group was lying on the same side, forcing phenyl rotation, and weakening the edge-to-face interaction with W1080 (**Fig 4cd, S7**). This preferred geometry, together with the fluorine-occupied hydrophobic pocket, provided a clear direction for analog design to improve potency.

In parallel, determination of absolute configuration using vibrational circular dichroism (VCD)^30^ of the enantiomerically pure compounds confirmed that the (R)-enantiomer was the active form (**Fig S8**), in agreement with AS-MS data, the map fitting scores, and FEP analysis.

### Structure activity relationship (SAR) insights for AS-MS Hit 1

Using structure-based drug design and virtual screening, we selected analogs that featured replacement or modification of the three distinct moieties (difluorocyclopropane, 1-methyl-1N-indazole, and N-methylaniline). **3**-**18** have modifications at one moiety and **19**-**23** at two moieties (**Fig 5**).

**Figure 5.**
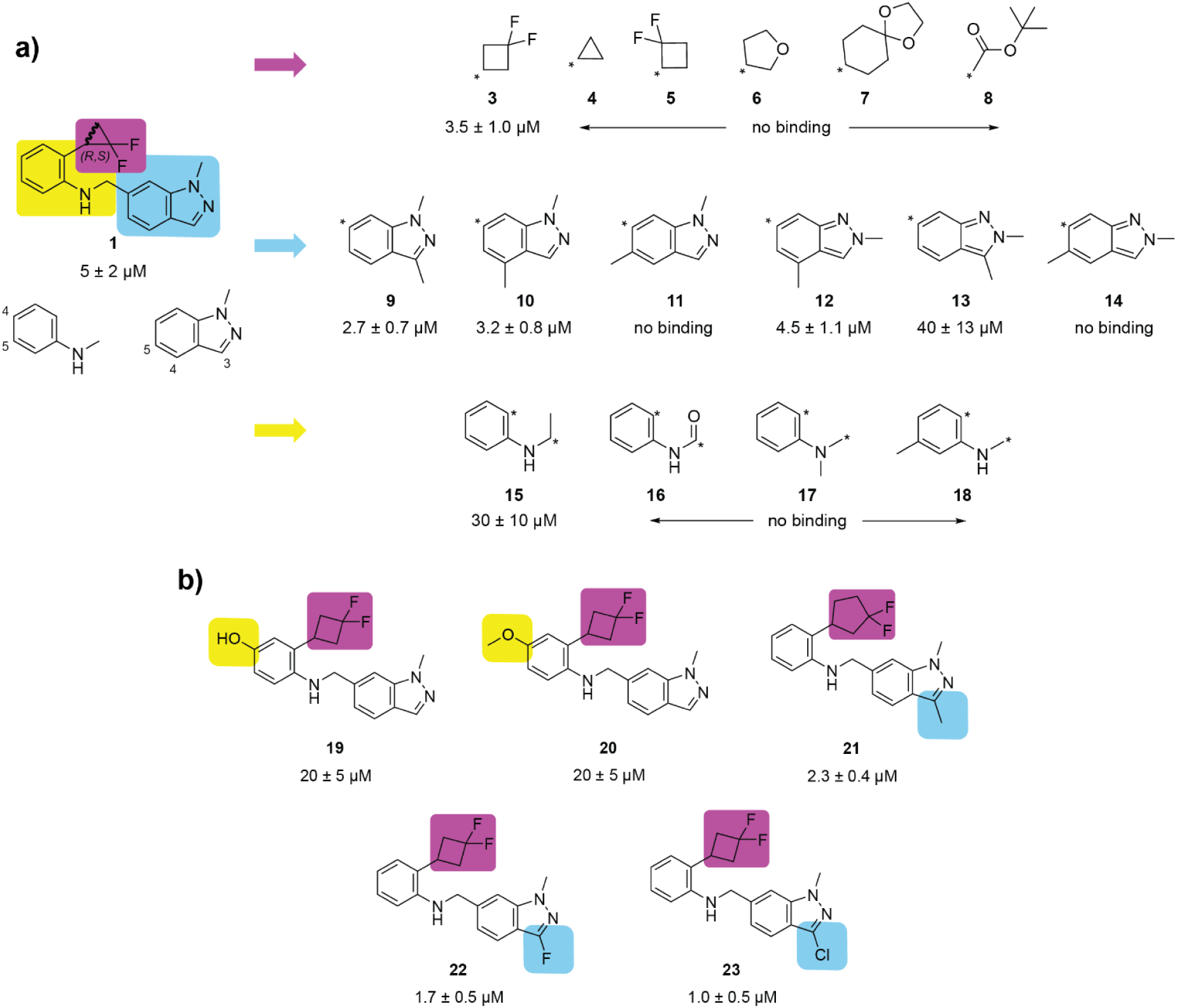
Structure and activity of analogs. AS-MS Hit 1. has three distinct moieties: difluorocyclopropane, 1-methyl-1H-indazole (core), and N-methylaniline. Analogs of **AS-MS Hit 1** that have **a)** one modification in one of the moieties, and **b)** modifications in two moieties. The K_D_ values were determined by SPR (N≥4, see **Table S1** for compiled SPR data). The SMILES and catalog numbers can be found in the Supporting Information.

Difluorocyclopropane could be replaced by 3,3-difluorocyclobutane (**3**), but cyclopropane (**4**), or 2,2-difluorocyclobutane (**5**) disrupted binding and unrelated, bulky moieties tetrahydrofuran (**6**), 1,4-dioxaspiro[4.5]decane (**7**), or tert-butyl acetate (**8**) also abrogated binding. We next explored adding a methyl group to various positions on the indazole, as well as regioisomers. Both regioisomers tolerated an additional methyl group at C4 (**10** and **12**) but not at C5 (**11** and **14**). C5 methylation may introduce a steric clash with the N-methylaniline, preventing a favourable binding pose. Methylation at C3 was well-tolerated in 1-methyl-1H- (**9**) but much less so for 2-methyl-2H-indazole (**13**). These trends suggest that the nitrogen substitution pattern of the indazole ring influences the binding pose and the π-π interaction with W1080. Modifications to the methylene carbon (**15**, **16**) or nitrogen (**17**) were not tolerated, suggesting the right angle made at the N-methyl of the N-methylaniline (**Fig 4d**) is required.

Unexpectedly, methylation at position 5 of N-methylaniline (**18**) disrupted binding. Inspection of the cryo-EM structure suggests that the methyl group might clash with H132 on HAP40 (**Fig 4d**).

We also investigated combining modifications. **19** and **20** combined 3,3-difluorocyclobutane (**3** in **Fig 5a**) with hydroxyl (**19**) and methoxy (**20**) on C4 of N-methylaniline (**Fig 5b**). **21** combined 3,3-difluorocyclopentane with methyl on C3 of 1-methyl-1H-indazole (**9** in **Fig 5a**). These modifications were well-tolerated but did not improve affinity. Combining the 3,3-difluorocyclobutane (**3** in **Fig 5a**) with fluorine (**22**) or chlorine (**23**) on C3 of 1-methyl-1H-indazole improved binding (**Fig 5b**). The halogenation may strengthen π-stacking with W1080 and/or introduce favourable interactions with nearby hydrophobic residues such as L88 on HAP40 (**Fig 4d**).

### Virtual screening pipeline to further expand SAR

We developed a structure-based virtual screening pipeline based on docking and molecular dynamics (MD) simulations in the internal coordinate space with ICM (Molsoft, San Diego). To test the computational workflow, we used it to retrospectively rank 44 analogs (comprising 11 binders and 33 non-binders), leading to five binders in the top 10 (**Fig 6ab**). For prospective selection of analogs, we first conducted substructure and similarity searches of **AS-MS Hit 1** against more than 10 billion compounds in the Enamine REAL library with SpaceMACS (BioSolveIT), yielding ∼2,500 analogs that were docked with ICM. The top 400 scoring compounds were advanced to MD, leading to 90 stable protein-ligand complexes, 72 of which were tested experimentally. While low solubility prevented conclusive assessment of multiple candidates, compounds **24** and **25** were clearly confirmed by SPR (**Fig 6cd**). The pyridine ring of **24** occupies a cavity that was not exploited by previous ligands, highlighting a potential vector for further optimization.

**Figure 6.**
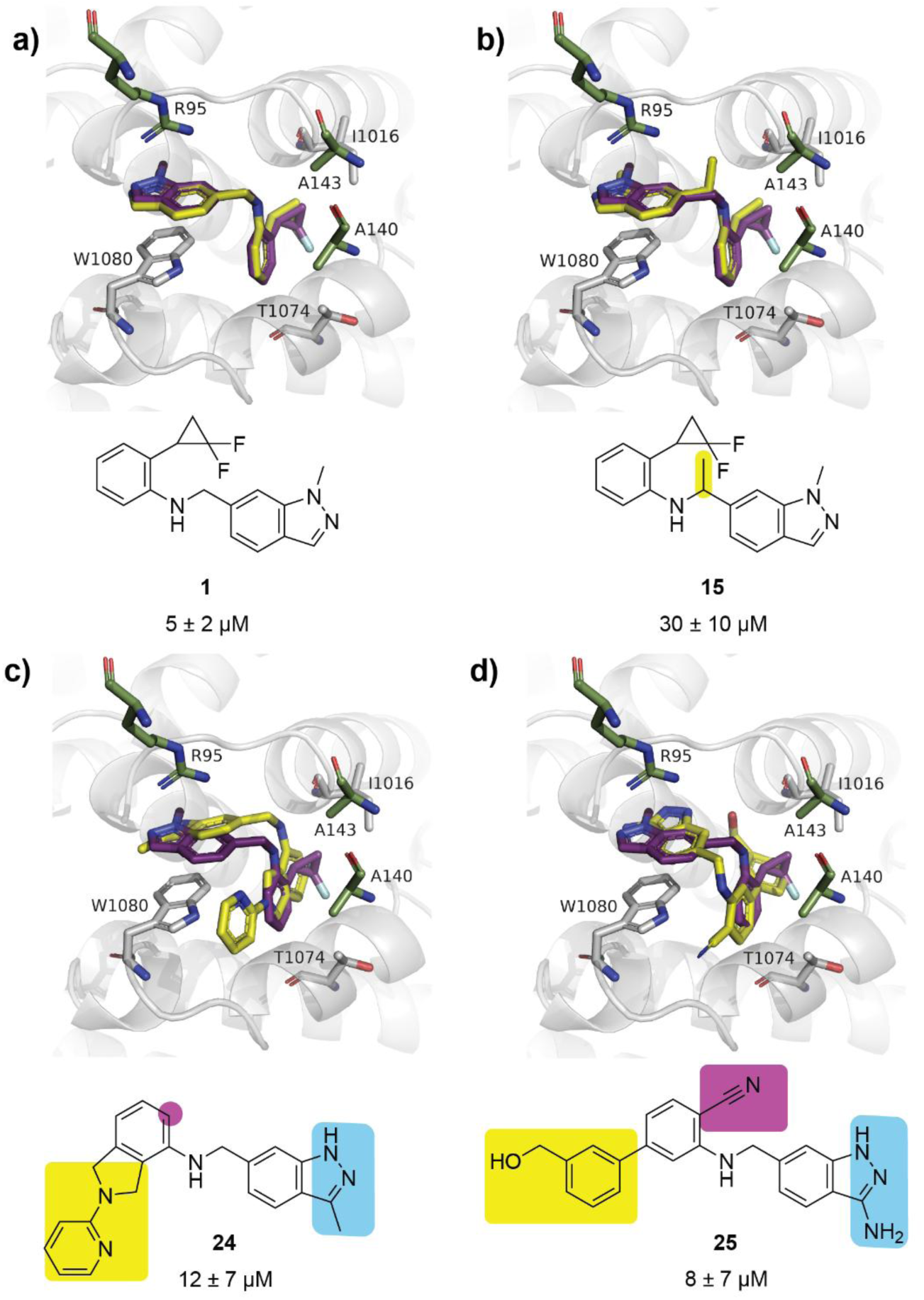
Docked poses of analogs from retrospective and prospective virtual screening. **a)** Redocked pose of **AS-MS Hit 1**. **b)** Docked pose for **15**, an active compound of the retrospective virtual screen. Docked poses for **c) 24** and **d) 25**, both active compounds from the prospective virtual screen. All docked poses are shown in yellow overlaid with the experimental pose of **AS-MS Hit 1** in purple. K_D_ values were determined by SPR (N≥3).

## Conclusions

We identify, biophysically characterise, and structurally validate the first small molecule ligands for full-length HTT in complex with HAP40. Our data reveal a previously unexploited, druggable binding pocket and demonstrate that ligand engagement is independent of polyglutamine expansion. Preliminary SAR exploration reveals potential avenues for further optimization. These results establish a reproducible framework for ligand discovery against HTT and provide a robust chemical starting point for probing HTT structure and function.

### Significance

We have identified binders for HTT that can underpin the development of HTT-directed probes, degraders, and imaging agents, opening a new path for therapeutic innovation in HD.

## Experimental Section

### Protein Expression and Purification

HTT proteins were purified as described previously.^21,22^ Briefly, HTT bearing polyQ tracts of Q23 or Q54 in either its apo or HAP40-bound forms, as well as structure-rationalized HTT domains (CTD res. V2095-V3138, NTD res. T97-M2069) were expressed by baculovirus-mediated transduction in sf9 insect cells. Purified proteins were obtained using FLAG affinity chromatography and subsequent size-exclusion chromatography. C-terminally biotinylated HTT (apo and in complex with HAP40) used in SPR was generated by cloning in a C-terminal Avi-tag and followed by co-expression with BirA and supplementation with exogenous biotin.

### Affinity Selection Mass Spectrometry (AS-MS)

Hit identification by AS-MS was carried out following the method described by Wang et al.^23^ Briefly, His-tagged HTT-HAP40 complex was immobilised on magnetic Ni-beads along with a panel of unrelated proteins to serve as a counter screen. The complete panel was incubated with pools of ∼500 compounds per pool, screening a total library of 15K compounds. The immobilised proteins were washed and bound ligands released by protein denaturation prior to identification by liquid chromatography-mass spectrometry (LC-MS) analysis. The relative enrichment of each compound was determined by comparing the mass spectral intensities of each compound for HTT-HAP40 compared to the other proteins in the panel. Compounds showing selective binding, defined as a >∼5-fold enrichment over other proteins in the panel, were prioritized as potential hits for downstream confirmation studies. **AS-MS Hit 1** was also verified for enantiomeric selectivity, in which both the chemical library stock and the eluates extracted from the initial AS-MS screen were separated using a CHIRALPAK IBN (250 × 2.1 mm) column from DAICEL Chemical Industries, LTD. LC gradient method 2 was used, as described by Wang et al.^23^

### Compound Purity

All reported compounds have a purity of at least 89% at 254 nm (**Fig S9**). The purity of the compounds was determined using a liquid chromatograph equipped with a diode array and a mass spectrometer detector (Waters Acquity SQD mass spectrometer).

### Surface plasmon resonance (SPR)

0.7-0.8 mg/mL HTTQ23-HAP40 (C-terminal biotin) was immobilized on a Streptavidin chip in SPR Running Buffer (10 mM HEPES pH 7.4, 150 mM NaCl, 1 mM EDTA, 0.005% Tween-20 (v/v), 0.2% PEG3350 (w/v), 4% DMSO (v/v)) to a response of 7000-9000 RU. The HTTQ23-HAP40 subdomains CTD-HAP40 and NTD (both C-terminal biotin) were immobilized at 0.75 mg/ml to ∼11000-15000 RU and ∼10000-12000 RU, respectively. WDR55 (N-terminal biotin) was immobilized at 0.2 mg/ml to ∼6000 RU. Multi Cycle Kinetics were performed for 6-8 point titrations of compounds serially diluted 2 or 3-fold. A 1:1 binding model was used to calculate the kinetic dissociation coefficient (K_D_) and the fit of the theoretical curve was determined by Chi^2^ (RU^2^). The %R_max_ was calculated by normalizing the observed maximum response of the curve (R_max_) to the theoretical maximum response given 1:1 binding (R_max theoretical_) which can be calculated with the following:

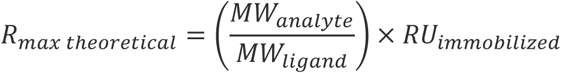

### Hydrogen-Deuterium Exchange Mass Spectrometry (HDX-MS)

HDX-MS was performed as previously described using the PAL3 Autosampler (CTC Analytics AG) coupled to an M-Class UPLC (Waters Corp., U.K.) and SELECT SERIES Cyclic IMS Mass Spectrometer (Waters Corp., U.K.) in HDMSe mode.^31^ Briefly, samples of 15 µM HTTQ23-HAP40 or HTTQ23-HAP40 + **AS-MS Hit 1** (Internal ChemiReg ID XS837367a) (1:5) were labeled (10 mM Phosphate Buffer pD 7.4, 150 mM NaCl) at 20 °C for 0, 0.25, 1, and 10 mins. The HDX reaction was then ‘harsh’ quenched (7.5 M Guanidine-HCl, 0.5 M TCEP, 100 mM phosphate buffer pH 2.5) at 0 °C for 2 min followed by dilution with an equal volume of quench dilution buffer (100 mM Phosphate Buffer, pH 2.5). The sample was digested (Nepenthesin2-Pepsin 1:1, Affipro), followed by desalting (ACQUITY UPLC BEH C18 VanGuard Pre-column, Waters Corp., U.K.), and reverse-phase separation (ACQUITY UPLC BEH C18 Column, Waters Corp., U.K.). Data processing was conducted using ProteinLynx Global Server 3.0.3. and DynamX 3.0 (Waters Corp., U.K.), and visualization was done using the AlphaFold3^32^ HTQ23-HAP40 model in PyMol 2.5.0 (Schrodinger, LLC). For a cumulative ΔHDX-MS peptide signal to be considered statistically significant, it must have exceeded the cumulative propagated error (3σ, n ≥ 2). For peptide sequence coverage, see **Figure S3.**

### Cryogenic electron microscopy (Cryo-EM)

#### Cryo-EM sample preparation and data acquisition

Four microliters of HTT-HAP40 (0.4 mg/ml) with 25 µM XS837367a in 20 mM HEPES pH 7.4, 150 mM NaCl, 1 mM TCEP, 0.025% (w/v) CHAPS was adsorbed onto lightly glow-discharged (10 s, 15 mA) suspended monolayer graphene grids (Graphenea) for 60 s. Grids were then blotted with filter paper for 1 s at 100 % humidity, 11 °C and frozen in liquid ethane using a Vitrobot Mark IV (Thermo Fisher Scientific).

Movies were collected in counted mode in Electron Event Representation (EER) format on a CFEG-equipped Titan Krios G4 (Thermo Fisher Scientific) operating at 300 kV with a Selectris X imaging filter (Thermo Fisher Scientific) with slit width of 10 eV, at 165,000x magnification on a Falcon 4i direct detection camera (Thermo Fisher Scientific) corresponding to a calibrated pixel size of 0.732 Å. Movies were collected at a total dose of 51.1 e^-^/Å^2^ fractionated to ∼ 1 e^-^/Å^2^ per fraction. In total, 10,990 movies were collected with an untilted stage, and 11,935 movies were collected with a stage tilt of 30° to improve particle angular distributions.

#### Cryo-EM data processing, model building, and refinement

Patched motion correction, CTF parameter estimation, particle picking, extraction, and initial 2D classification were performed in SIMPLE 3.0.^33^ All downstream processing was carried out in cryoSPARC or RELION 3.1,^34,35^ using the csparc2star.py script within UCSF pyem (Asarnow, D., Palovcak, E., Cheng, Y. UCSF pyem v0.5. (*Zenodo:2019*)) to convert between formats. Global resolution was estimated from gold-standard Fourier shell correlations (FSCs) using the 0.143 criterion and local resolution estimation was calculated within cryoSPARC.

The cryo-EM processing workflow is outlined in **Figure S6**. Briefly, downsampled particles (1.464 Å / pixel, 256 x 256 pixel box size) from the untilted or tilted datasets were each independently subjected to a round of reference-free 2D classification (k=200 each) using a 180 Å soft circular mask within cryoSPARC. 1,549,562 particles were selected from the untilted dataset after 2D classification and subjected to multi-class *ab initio* reconstruction using a maximum resolution cutoff of 6 Å, generating four volumes. These volumes were subsequently used as reference volumes (after lowpass filtering to 20 Å) for a heterogeneous refinement against the full 2D-selected untilted particle stack. Particles (1,135,563) from the most populated and structured class were selected and non-uniform refined against their corresponding volume lowpass-filtered to 30 Å, generating a Nyquist-limited 3.0 Å map. For the tilted particle set, 1,273,790 particles were selected after 2D classification and subjected to heterogeneous refinement against the same 20 Å lowpass-filtered references used previously for the untilted dataset. Particles (993,717) from the most populated and structured class were selected and non-uniform refined against their corresponding volume lowpass-filtered to 30 Å, also generating a Nyquist-limited 3.0 Å map. Bayesian polishing was then performed independently in RELION for both particle stacks, reverting to the nominal movie pixel size of 0.732 Å and a box size of 512 x 512 pixels. Further 2D classification and non-uniform refinement (with local CTF refinement) of each particle set resulted in a 2.2 Å map from 1,114,910 untilted particles and a 2.3 Å map from 943,456 tilted particles, and a 2.1 Å map when the datasets were combined and non-uniform refined. Particles from the combined refinement were further subjected to 3D classification within cryoSPARC (k=8) using a soft dilated mask encompassing the entire volume and a resolution filter of 3 Å. The most populated class, containing 387,879 particles, demonstrated clear density for N-terminal helices of HAP40, whereas two other classes (containing 287,941 and 305,684 particles) lacked this density; but all three classes contained equivalent density for **AS-MS Hit 1**. Non-uniform refinement of the most populated class (conformer 1) resulted in a 2.3 Å map, and non-uniform refinement of the other two classes (conformers 2 and 3) resulted in 2.4 Å maps.^33–36^

### Vibrational circular dichroism (VCD)

#### VCD Measurements

The CDCl_3_ solvent (Cambridge Isotope Labs Silver Foil) was run through a small plug of activated basic alumina immediately before use. To a small vial containing 6 mg of each enantiomer of **AS-MS Hit 1**, 110 uL of CDCl_3_ was added. The resulting solution was transferred to a liquid IR cell (BaF_2_, 100 μm cell path) and placed in the measurement chamber. The instrumentation is at BioTools, Inc. (Jupiter, FL) ChiralIR 2X DualPEM™ FT-VCD spectrometer, set to 4 cm^-1^ resolution, with PEM (both 1 and 2) maximum frequency set to 1400 cm^-1^. The sample was then measured for 8 hours in one hour blocks. The IR data from the first block was solvent and water vapor subtracted, then offset to zero at 2000 cm^-1^. The VCD data blocks were averaged, and the baseline corrected using enantiomer subtraction ((E1 – E2) / 2). Finally the VCD spectrum was offset to zero at 2000 cm^-1^. The VCD noise data was block averaged and used without further processing.

### VCD Calculations

The (R)-enantiomer was constructed using ComputeVOA (BioTools, Jupiter, FL). A thorough conformational search was performed at the MM level using the MMF94 force field in a 7 kcal/mol energy window. All 58 conformers found were subjected to DFT level optimization and frequency calculation with Gaussian ’09^37^ (Wallingford, CT)**^VCD^** using the B3LYP / 6-31G(d) and B3PW91 / 6-31G(d) methods using the implicit solvation CPCM (chloroform) method. The resulting lowest energy unique conformations (∼20 for each method) were reoptimized using the B3LYP / cc-pVTZ and B3PW91 / cc-pVTZ methods respectively, and the IR and VCD frequencies recalculated at this level. The resulting spectra from all methods were Boltzmann averaged (using both Free Energy and Electronic Energy), plotted at 5 cm^-1^ resolution and then x-axis scaled (range of 0.968 to 0.985 - values obtained using CompareVOA and varied with basis set and functional) for comparison to the experimental IR and VCD spectra. The two different weighing methods (Free Energy and Electronic Energy) gave consistent results for stereochemistry, with the best results coming from the electronic energy weighted average of the B3PW91 / cc-pVTZ (CPCM – chloroform) DFT method. The resulting DFT calculated IR and VCD spectra of the (R)-enantiomer matched that of **AS-MS Hit 1**, and gave high similarity values for both IR and VCD and a 100% confidence level match in CompareVOA (BioTools Inc. Jupiter, FL).^38,39^ All plots and comparisons are using this data.

### In silico Studies

#### Free-energy perturbation (FEP)

FEP calculations were performed using FEP+ and the OPLS4 force field.^28,29,40^ The HTT-HAP40 structural models were prepared by the Protein Preparation Workflow^41,42^ using the initial model for cryo-EM structure (PDB 9YR6). Ligands were prepared by Ligprep.^43^ FEP+ simulations were performed for 15 ns using 20 lambda windows.^28,29^

#### *In silico*-based virtual screening: Docking

The HTT-HAP40-Compound 1 cryo-EM structure was first subjected to the default protein-preparation module implemented in ICM (Molsoft, San Diego). Briefly, missing sidechains and hydrogen atoms were added and energy-minimized; hydrogen-bond networks were further optimized by scanning terminal rotameric states of asparagine and glutamine and by sampling the tautomeric states of histidine sidechains. The grid representation of the binding pocket (rigid) was used to docked compounds (flexible). Generated docked poses were assigned a physics-based score and a radial and topological convolutional neural network (RTCNN) score. As both scores are complementary, they were added to rank-order docked poses with a cut-off for summed scores greater than -41 kcal/mol.

#### *In silico*-based virtual screening: Molecular dynamics

The molecular dynamics module of ICM (Molsoft) based on the OpenMM forcefield was deployed to identify the most stable binding poses of each compound. A box surrounding the complex was flooded with water and MD was run for 5 ns. Molecular drift and fit across 20 frames evenly spaced along the MD trajectories were analyzed using the provided root mean square deviation (RMSD) and “RTCNN_mean” metrics respectively.

## Ancillary Information

### Supporting Information Availability

The Supporting Information comprises:

SPR data, HDX-MS peptide sequence coverage, differential data, and kinetic plots, cryo-EM data collection and refinement statistics and workflow, binding poses of both enantiomers of **1**, VCD of (R)-**1**, and HPLC purity analysis for **1** - **25** (PDF)

Any additional information is available from the lead author upon request.

## Author Information

### Corresponding authors

**Rachel J. Harding** − Structural Genomics Consortium, University of Toronto, Toronto, Ontario M5G 1L7, Canada; Department of Pharmacology and Toxicology, University of Toronto, Toronto, Ontario M5S 1A8, Canada; Princess Margaret Cancer Centre, University Health Network, Toronto, ON M5G 2M9, Canada; Leslie Dan Faculty of Pharmacy, University of Toronto, Toronto, ON M5S 3M2, Canada; orcid.org/0000-0002-1134-391X;

## Author Contributions

The manuscript was written by R.J.H., R.L., S.A., E.W., S.B., C.V., and S.F. and revised by M.S. All authors have given approval to the final version of the manuscript. R.C. purified the proteins and J.S. and X.W. performed the AS-MS. All compounds were purchased from Enamine or ChemDiv and A.B. prepared the compounds. R.L. performed the SPR, E.W. performed the HDX-MS, J.C.D. collected the cryo-EM data and generated the maps, and S.B. refined the cryo-EM model. S.F. conducted the FEP calculations, S.A. led SAR investigations and compound analog design, and C.V.S. created and conducted the virtual screening workflow. R.J.H., M.S., J.D.M, Y.Z., R.K., D.W., S.M.L., A.M.E., C.H.A, H.P., and L.H. advised throughout the project.

## Declaration of Interests

The authors declare no competing financial interest.

## Acknowledgements

The Structural Genomics Consortium is a registered charity (no: 1097737) that receives funds from Bayer AG, Boehringer Ingelheim, Bristol Myers Squibb, Genentech, EU/EFPIA/OICR/McGill/KTH/Diamond Innovative Medicines Initiative 2 Joint Undertaking [EUbOPEN grant 875510], Janssen, Pfizer, and Takeda. The Harding lab is supported by funding from CIHR (Funding reference number: 198025), NSERC (RGPIN-2024-05769), CFI, the Hereditary Disease Foundation, Schrödinger LLC, the Connaught Fund, and Conscience – DMOS. We thank the Structural & Biophysical Core (SBC) Facility at The Hospital for Sick Children (SickKids) for access to instrumentation and technical support. We thank BioTools for performing the vibrational circular dichroism (VCD) experiments.

## Notes

### Competing Interest Statement

The authors have declared no competing interest.

