## Supplementary Information for "Identification and Characterization of a Small Molecule Ligand for the Huntingtin-HAP40 Complex"

[1] Structural Genomics Consortium, University of Toronto, 101 College St, Toronto, ON M5G 1L7, Canada
[2] Department of Pharmacology and Toxicology, University of Toronto, Toronto, ON M5S 1A8, Canada
[3] Department of Chemistry, York University, Toronto, ON M3J 1P3, Canada
[4] Center for Structural Biology, Center for Cancer Research, National Cancer Institute, Frederick, MD, USA
[5] Schrödinger, Inc., 1540 Broadway, 24th Floor, New York, New York 10036, United States
[6] Department of Chemistry, University of Toronto, ON M5S 3H6, Canada
[7] Department of Molecular Genetics, Faculty of Medicine, University of Toronto, Toronto, ON M5S 1A8, Canada
[8] Department of Medical Biophysics, University of Toronto, Toronto, ON, M5G1L7, Canada
[9] Princess Margaret Cancer Centre, University Health Network, Toronto, ON M5G 2M9, Canada
[10] Leslie Dan Faculty of Pharmacy, University of Toronto, Toronto, ON M5S 3M2, Canada

**Contents of Supporting Information**

**S2:** Table S1. Summary of SPR results for primary AS-MS hits and analogs
**S5:** Figure S1. Characterization of AS-MS Hit 1 binding to HTTQ54-HAP40 by SPR
**S6:** Figure S2. Characterization of binding for AS-MS Hit 2 against HTT-HAP40 subdomains and WDR55
**S7:** Figure S3. HTTQ23 and HAP40 peptide sequence coverage in HDX-MS
**S8:** Figure S4. Differential HDX-MS of HTTQ23-HAP40 + AS-MS Hit 1 (1:5) minus HTTQ23-HAP40
**S9:** Figure S5. Deuterium uptake kinetic plots of exemplary, a) HTT peptides and, b) HAP40 peptides
**S10:** Figure S6. Cryo-EM processing workflow and map quality metrics for HTT-HAP40 with bound AS-MS Hit 1
**S12:** Table S2. Cryo-EM data collection, refinement and validation statistics
**S13:** Figure S7. Cryo-EM binding pose interactions of AS-MS Hit 1 enantiomers
**S14:** Figure S8. Verification of AS-MS Hit 1 Peak 2 as the (R)-enantiomer by vibrational circular dichroism (VCD) and infrared spectroscopy (IR)
**S15:** HPLC traces of compounds 1 - 25

**Table S1. Summary of SPR results for primary AS-MS hits and analogs.**

| Compound | Protein Construct | $K_D$ (μM)[a] | $\%R_{max}$[a] | Affinity $Chi^2$ ($RU^2$)[a] |
|---|---|---|---|---|
| **1 AS-MS Hit 1** | HTTQ23-HAP40 | **5 ± 2** | **110% ± 12%** | **0.05 ± 0.07** |
| | CTD-HAP40 | Undetermined | Undetermined | Undetermined |
| | NTD | Undetermined | Undetermined | Undetermined |
| | WDR55 | Undetermined | Undetermined | Undetermined |
| | HTTQ54-HAP40 | **4 ± 1** | **120% ± 3%** | **0.2 ± 0.1** |
| **(S)-AS-MS Hit 1** | HTTQ23-HAP40 | Undetermined | Undetermined | Undetermined |
| **(R)-AS-MS Hit 1** | HTTQ23-HAP40 | **4 ± 2** | **100% ± 9%** | **0.05 ± 0.04** |
| **2 AS-MS Hit 2** | HTTQ23-HAP40 | **4 ± 2** | **230% ± 75%** | **0.82 ± 0.85** |
| | CTD-HAP40 | Undetermined | Undetermined | Undetermined |
| | NTD | Undetermined | Undetermined | Undetermined |
| | WDR55 | Undetermined | Undetermined | Undetermined |
| | HTTQ54-HAP40 | Undetermined | Undetermined | Undetermined |
| **3** | HTTQ23-HAP40 | **3.5 ± 1.0** | **110% ± 7%** | **0.06 ± 0.03** |
| **4** | HTTQ23-HAP40 | Undetermined | Undetermined | Undetermined |
| **5** | HTTQ23-HAP40 | Undetermined | Undetermined | Undetermined |
| **6** | HTTQ23-HAP40 | Undetermined | Undetermined | Undetermined |
| **7** | HTTQ23-HAP40 | Undetermined | Undetermined | Undetermined |
| **8** | HTTQ23-HAP40 | Undetermined | Undetermined | Undetermined |

| | | | | |
|---|---|---|---|---|
| **9** | HTTQ23-HAP40 | **2.7 ± 0.7** | **100% ± 7%** | **0.03 ± 0.02** |
| | HTTQ54-HAP40 | **2.6 ± 1.1** | **100% ± 8%** | **0.11 ± 0.08** |
| **10** | HTTQ23-HAP40 | **3.2 ± 0.8** | **100% ± 13%** | **0.03 ± 0.02** |
| **11** | HTTQ23-HAP40 | Undetermined | Undetermined | Undetermined |
| **12** | HTTQ23-HAP40 | **4.5 ± 1.1** | **100% ± 9%** | **0.04 ± 0.02** |
| **13** | HTTQ23-HAP40 | **40 ± 13** | **170% ± 23%** | **0.07 ± 0.03** |
| **14** | HTTQ23-HAP40 | Undetermined | Undetermined | Undetermined |
| **15** | HTTQ23-HAP40 | **30 ± 10** | **120% ± 22%** | **0.08 ± 0.05** |
| **16** | HTTQ23-HAP40 | Undetermined | Undetermined | Undetermined |
| **17** | HTTQ23-HAP40 | Undetermined | Undetermined | Undetermined |
| **18** | HTTQ23-HAP40 | Undetermined | Undetermined | Undetermined |
| **19** | HTTQ23-HAP40 | **20 ± 5** | **170% ± 27%** | **0.2 ± 0.2** |
| **20** | HTTQ23-HAP40 | **20 ± 5** | **140% ± 8%** | **0.2 ± 0.1** |
| **21** | HTTQ23-HAP40 | **2.3 ± 0.4** | **70% ± 10%** | **0.11 ± 0.06** |
| **22** | HTTQ23-HAP40 | **1.7 ± 0.5** | **60% ± 7%** | **0.06 ± 0.02** |
| **23** | HTTQ23-HAP40 | **1.0 ± 0.5** | **50% ± 12%** | **0.06 ± 0.03** |
| **24** | HTTQ23-HAP40 | **12 ± 7** | **20% ± 8%** | **0.02 ± 0.01** |
| **25** | HTTQ23-HAP40 | **8 ± 7** | **40% ± 16%** | **0.11 ± 0.03** |

[a]$K_D$ values are reported as mean ± standard deviation (N≥3). A 1:1 binding model was used to calculate the kinetic dissociation coefficient ($K_D$) and the fit of the theoretical curve was determined by $Chi^2$ ($RU^2$). The %$R_{max}$ was calculated by normalizing the observed maximum response of the curve ($R_{max}$)

to the theoretical maximum response given 1:1 binding ($R_{max\ theoretical}$) which can be calculated with the following:

$$R_{max\ theoretical} = \left(\frac{MW_{analyte}}{MW_{ligand}}\right) \times RU_{immobilized}$$

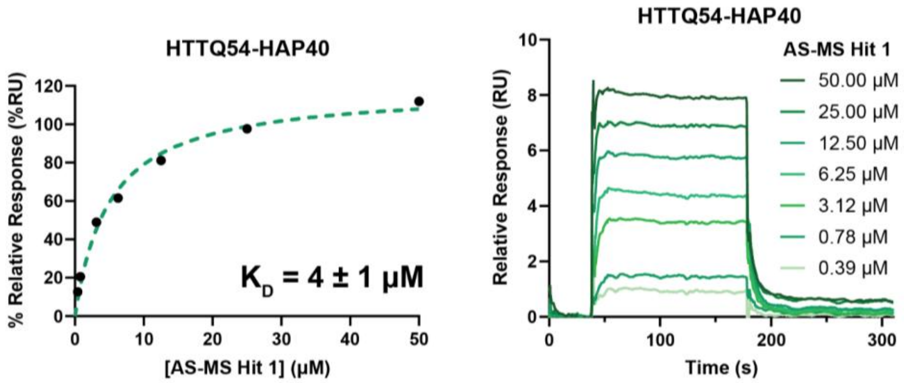


**Figure S1. Characterization of AS-MS Hit 1 binding to HTTQ54-HAP40 by SPR.** Steady state response (black circles) and a 1:1 binding model fit (dashed green line) were used. The raw sensorgram, affinity plot, and reported $K_D$ value (mean ± standard deviation) are representative of N=4.

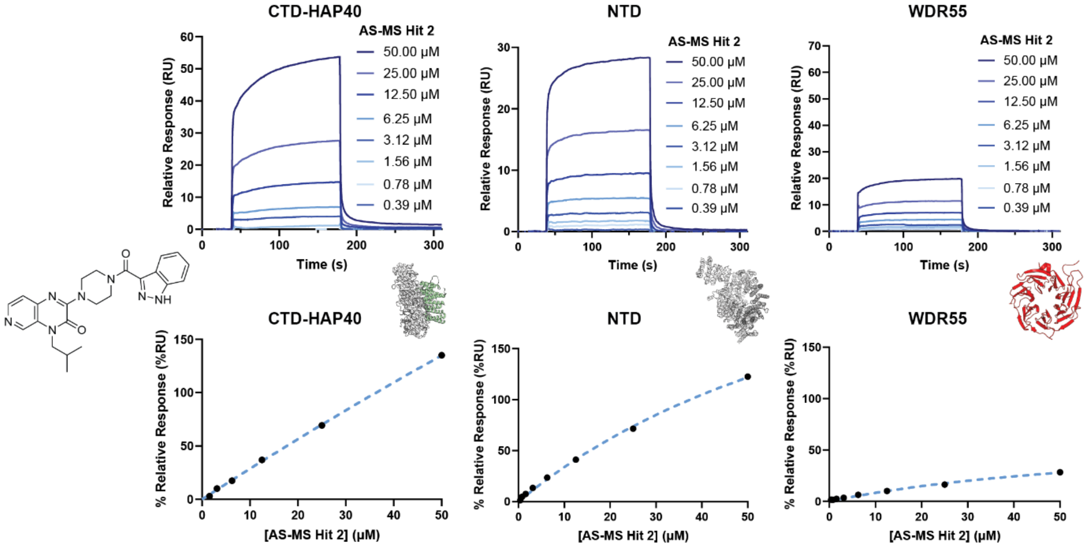


**Figure S2. Characterization of binding for AS-MS Hit 2 against HTT-HAP40 subdomains and WDR55.** Steady state response (black dots) is displayed with a 1:1 binding model (dashed line). Raw sensorgrams and affinity plots are representative of three replicates performed with the HTT subdomains and WDR55 (N≥3). $K_D$ is considered undetermined if the steady state response plot does not fully reach saturation and the $K_D$ is greater than half the maximum analyte concentration tested.

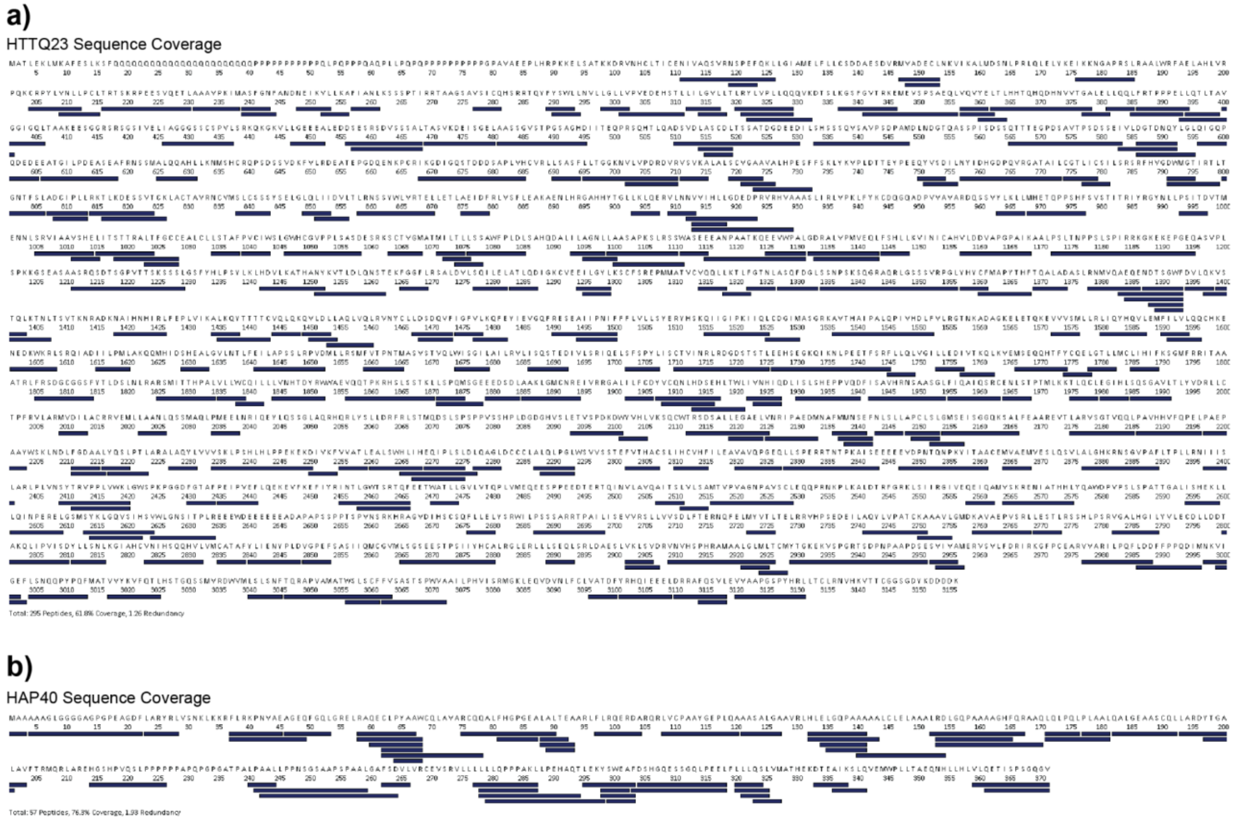


**Figure S3. HTTQ23 and HAP40 peptide sequence coverage in HDX-MS.** In this study, 295 HTTQ23 peptides yielded a sequence coverage of 61.8% and a peptide-per-residue redundancy of 1.26. Additionally, 57 HAP40 peptides yielded a sequence coverage of 76.3% and a peptide-per-residue redundancy of 1.93. These digests were obtained using an Affipro Nepenthesin2-Pepsin (1:1) protease column.

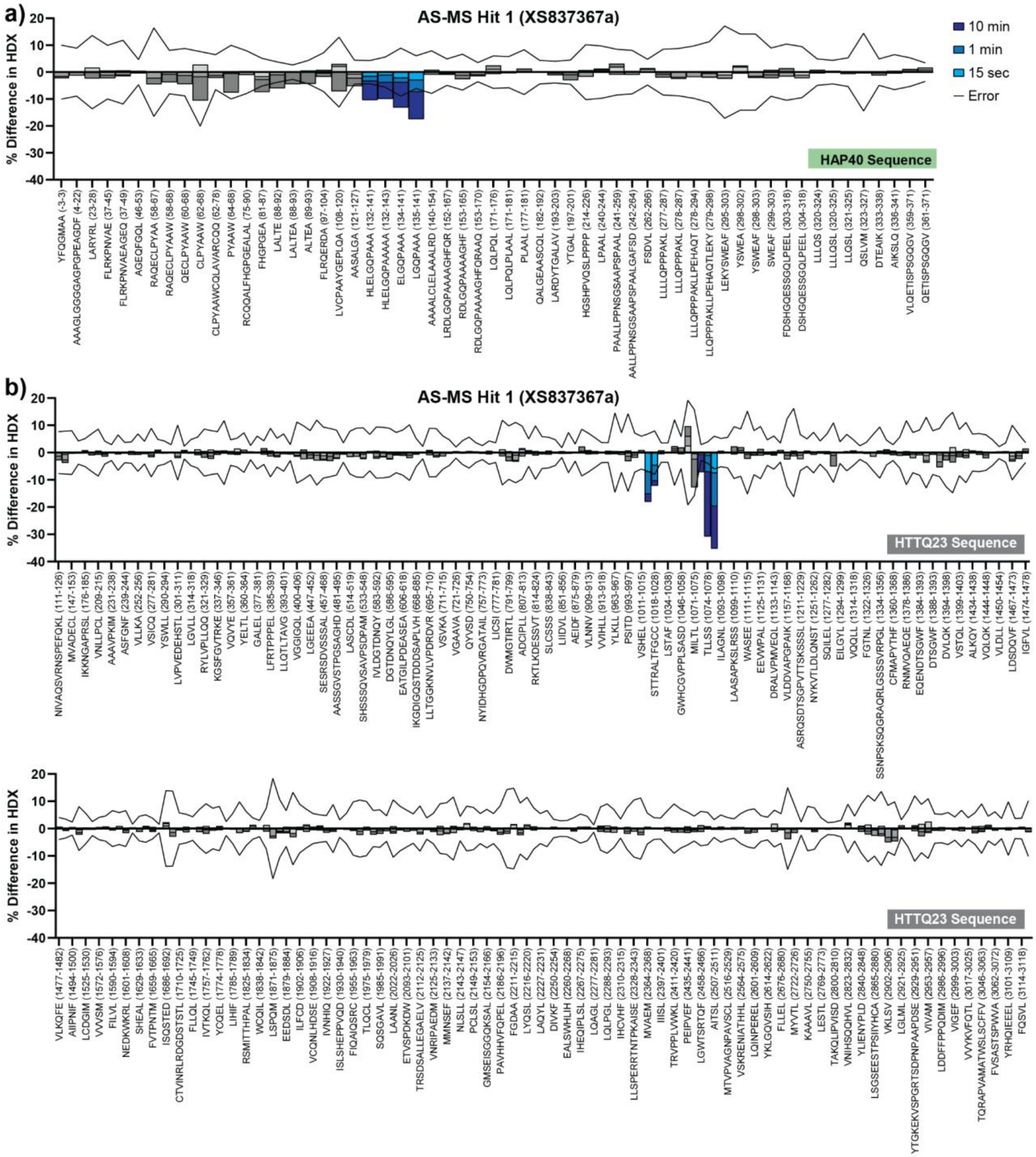


**Figure S4. Differential HDX-MS of HTTQ23-HAP40 + AS-MS Hit 1 (1:5) minus HTTQ23-HAP40.** Differential HDX is shown for timepoints of 15 sec (light blue), 1 min (blue), and 10 min (dark blue) for a) HAP40 peptides and b) HTTQ23 peptides. To be considered statistically significant, the cumulative differential HDX must exceed the error (black line) which represents triple the propagated standard deviation summed across all 3 timepoints (Wolf, E., et al. 2024, *Anal Chem*).

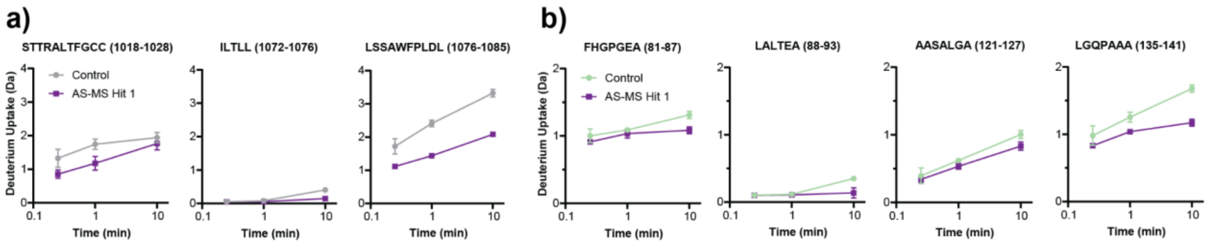


**Figure S5. Deuterium uptake kinetic plots of exemplary, a) HTT peptides and, b) HAP40 peptides.** The ligand-bound state (purple) is compared to the control/unbound state (grey-HTT, green-HAP40).

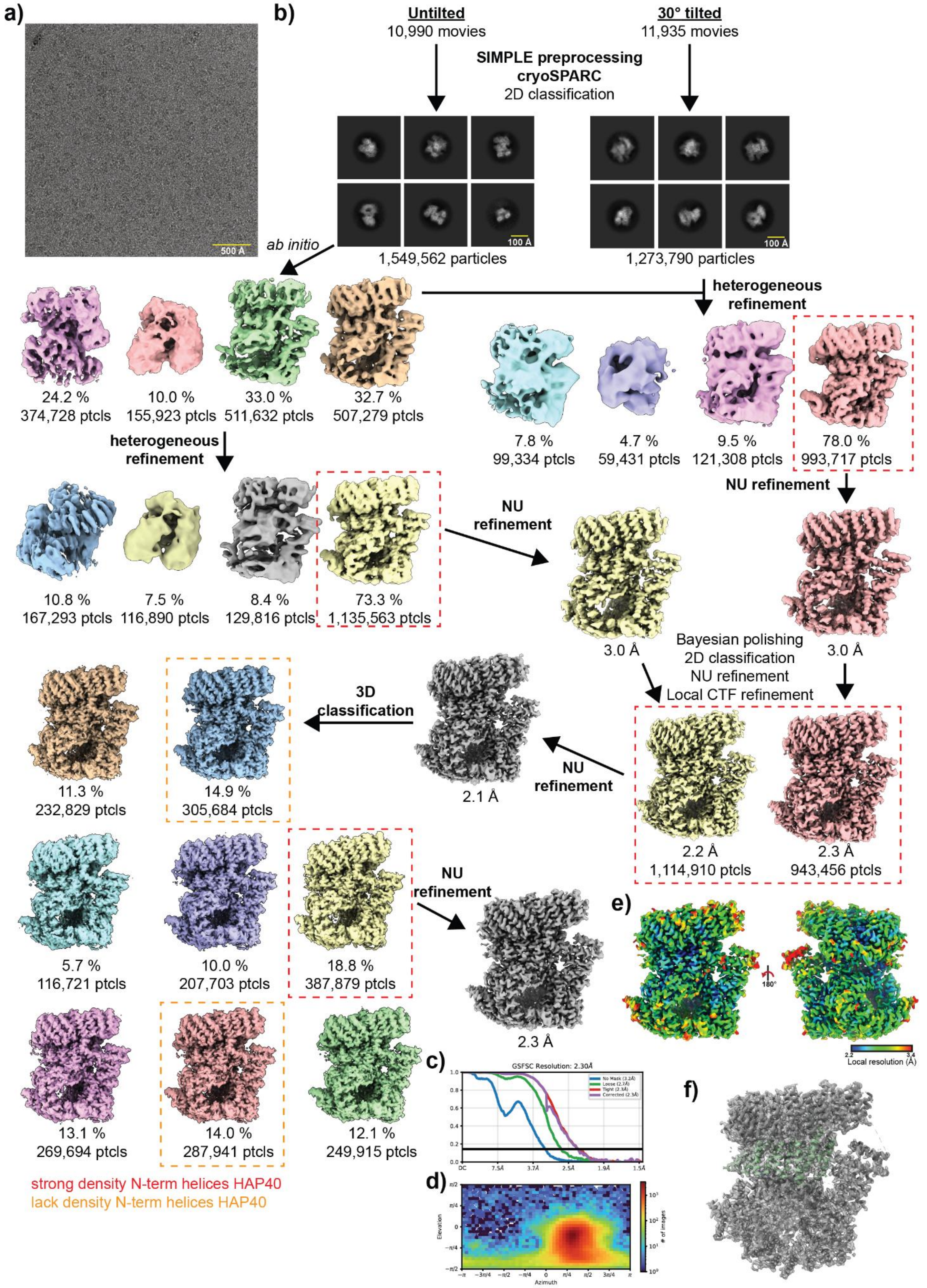


**Figure S6. Cryo-EM processing workflow and map quality metrics for HTT-HAP40 with bound AS-MS Hit 1. a)** Representative micrograph at -2 µm defocus. **b)** Image processing workflow. **c)** Global resolution estimate as determined by gold-standard Fourier shell correlations. **d)** Orientation distribution plots. **e)** Local resolution estimate of the volume. **f)** Final model of HTT-HAP40 in complex with **AS-MS Hit 1** with model/map overlay.

**Table S2. Cryo-EM data collection, refinement and validation statistics.**

| | HTT-HAP40-XS837367a (PDB 9YR6) (EMD-73361) |
|---|---|
| **Data collection and processing** | |
| Magnification | 165,000 |
| Voltage (kV) | 300 |
| Electron exposure (e–/Å$^2$) | 51.1 |
| Defocus range (μm) | -2.0 to -0.5 |
| Pixel size (Å) | 0.732 |
| Symmetry imposed | C1 |
| Initial particle images (no.) | 4,780,372 |
| Final particle images (no.) | 387,879 |
| Map resolution (Å) | 2.3 |
| FSC threshold | 0.143 |
| Map resolution range (Å) | 1.9-39 |
| **Refinement** | |
| Initial model used (PDB code) | 6X9O |
| Model resolution (Å) | 2.3 |
| FSC threshold | 0.143 |
| Map sharpening *B* factor (Å$^2$) | -55.5 |
| Model composition | |
| Non-hydrogen atoms | 19076 |
| Protein residues | 2709 |
| Ligands | 23 |
| *B* factors (Å$^2$) | |
| Protein | 30-134 |
| R.m.s.z[1] | |
| Bond lengths | 0.74 |
| Bond angles | 1.35 |
| Validation | |
| MolProbity score | 0.73 |
| Clashscore | 0.19 |
| Poor rotamers (%) | 1.4 |
| Ramachandran plot | |
| Favored (%) | 98 |
| Allowed (%) | 2 |
| Disallowed (%) | 0 |

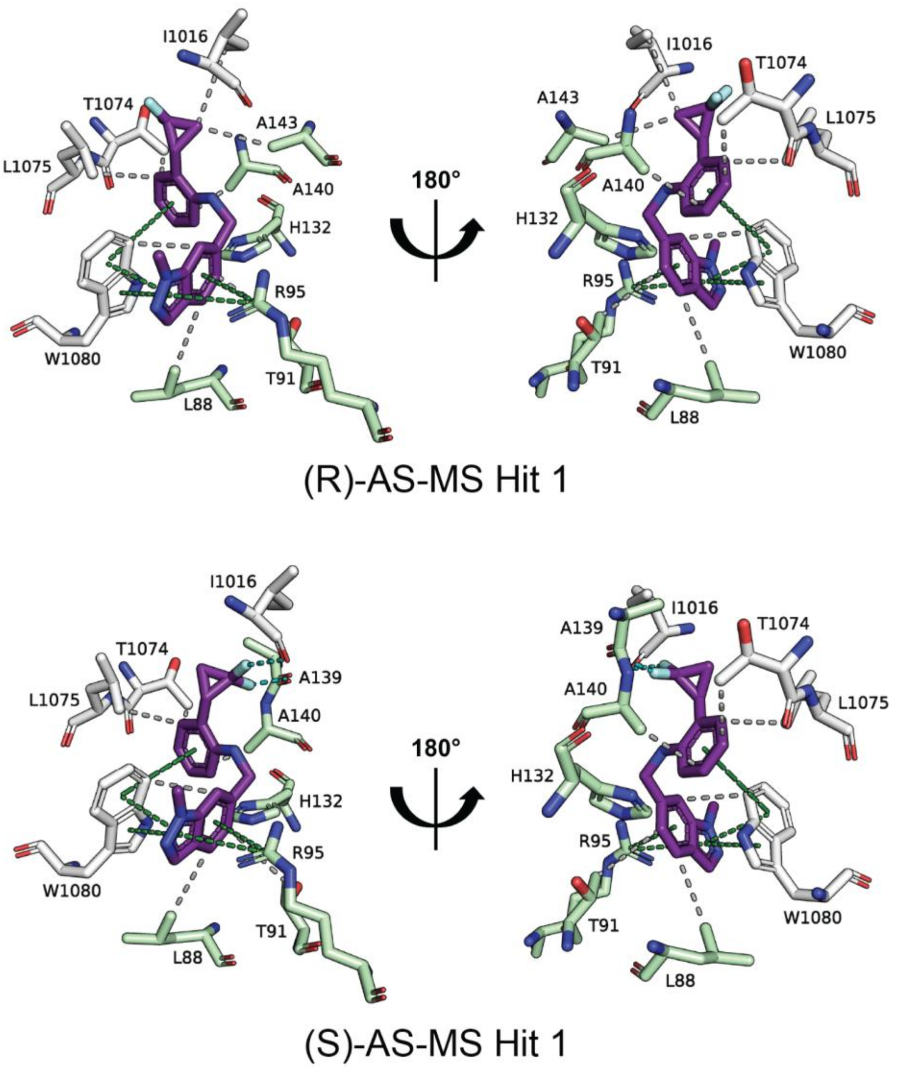


**Figure S7. Cryo-EM binding pose interactions of AS-MS Hit 1 enantiomers.** The (R)-enantiomer of **AS-MS Hit 1** (purple) binds using a combination of interactions shown in dashed lines: hydrophobic interactions (grey) and π-interactions (green). Residues from HTT are shown in white and those from HAP40 are shown in green. The (S)-enantiomer of **AS-MS Hit 1** binds with an interaction pattern very similar to that of the (R)-enantiomer, but with the addition of two halogen bonds (cyan dashed lines).

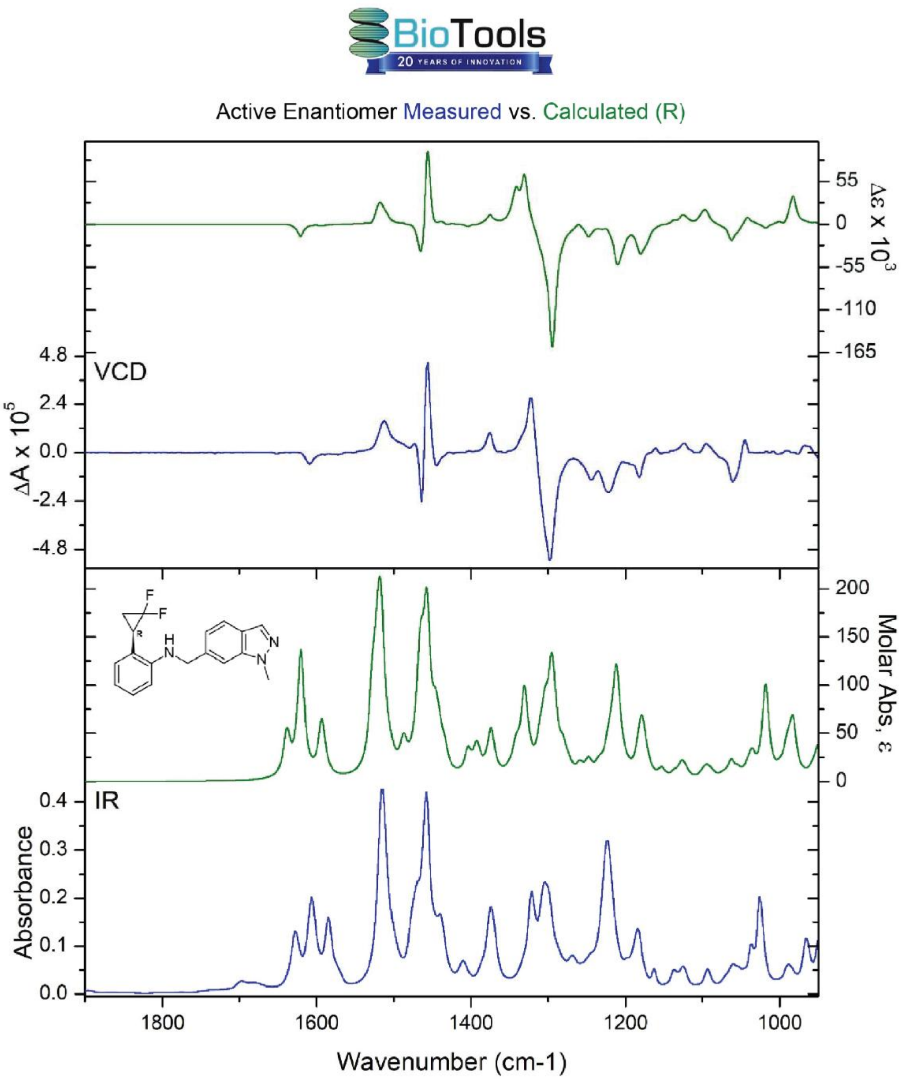


**Figure S8. Verification of AS-MS Hit 1 Peak 2 as the (R)-enantiomer by vibrational circular dichroism (VCD) and infrared spectroscopy (IR).** VCD (upper frame) and IR (lower frame) spectra measured for **AS-MS Hit 1** Peak 2 (left axes, blue spectra) compared with the Boltzmann-averaged spectra of the calculated conformations for the (R) configuration (right axes, green spectra).

### HPLC traces of compounds 1 - 25

*2-(2,2-difluorocyclopropyl)-N-((1-methyl-1H-indazol-6-yl)methyl)aniline, **1***

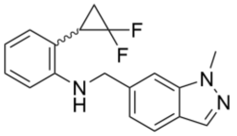

Chemical Formula: $C_{18}H_{17}F_2N_3$
Exact Mass: 313.14
Molecular Weight: 313.35

<u>**1** - Batch A:</u>

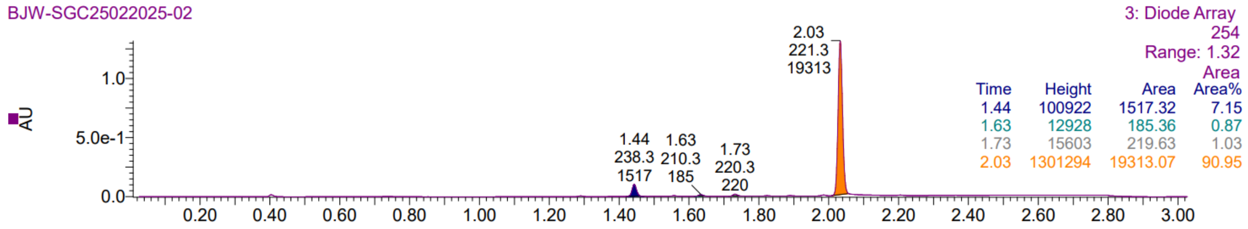


<u>**1** - Batch B:</u>

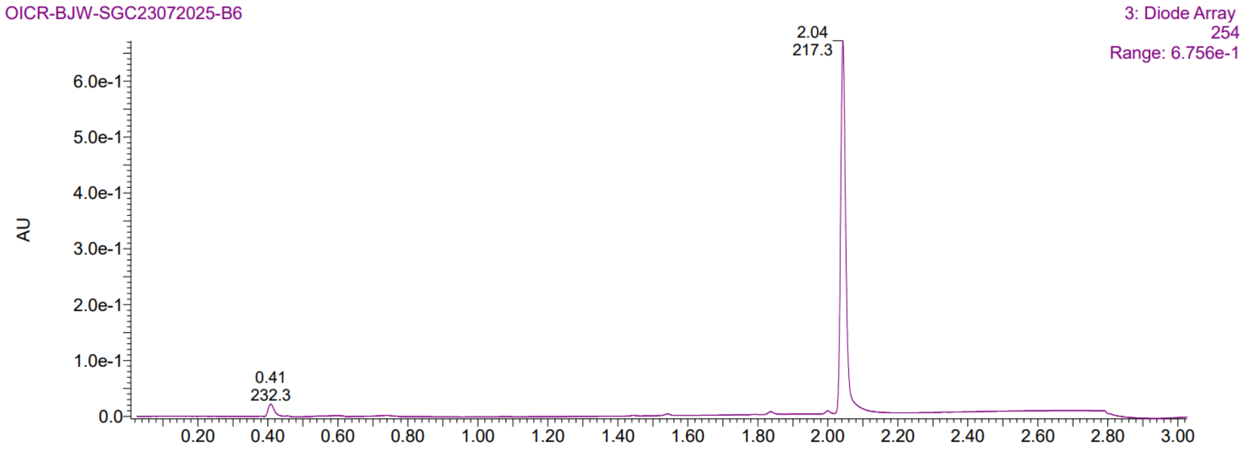


*(S)-2-(2,2-difluorocyclopropyl)-N-((1-methyl-1H-indazol-6-yl)methyl)aniline, **(S)-1***

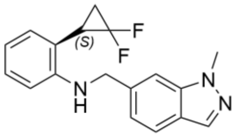

Chemical Formula: $C_{18}H_{17}F_2N_3$
Exact Mass: 313.14
Molecular Weight: 313.35

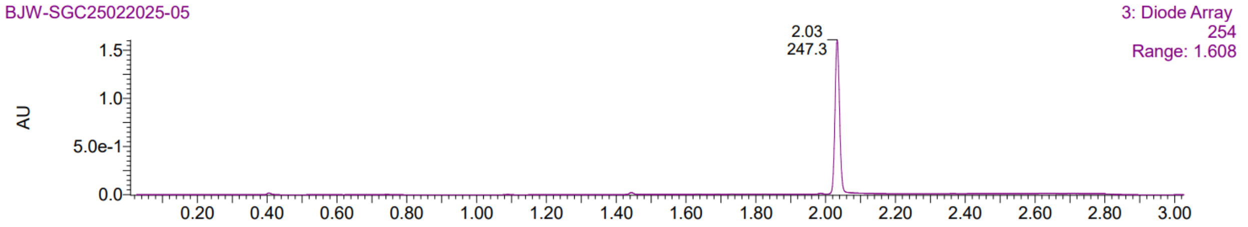

*(R)-2-(2,2-difluorocyclopropyl)-N-((1-methyl-1H-indazol-6-yl)methyl)aniline,* ***(R)-1***

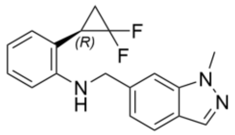


Chemical Formula: $C_{18}H_{17}F_2N_3$
Exact Mass: 313.14
Molecular Weight: 313.35

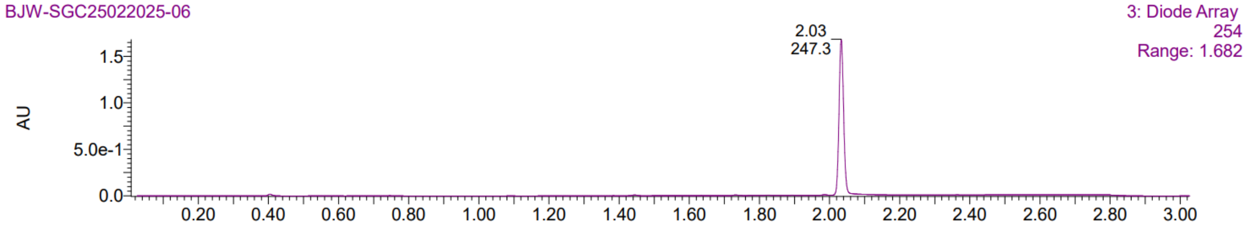


*2-(4-(1H-indazole-3-carbonyl)piperazin-1-yl)-4-isobutylpyrido[3,4-b]pyrazin-3(4H)-one,* ***2***

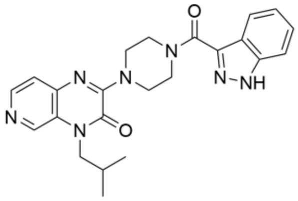


Chemical Formula: $C_{23}H_{25}N_7O_2$
Exact Mass: 431.21
Molecular Weight: 431.50

**2** - Batch A:

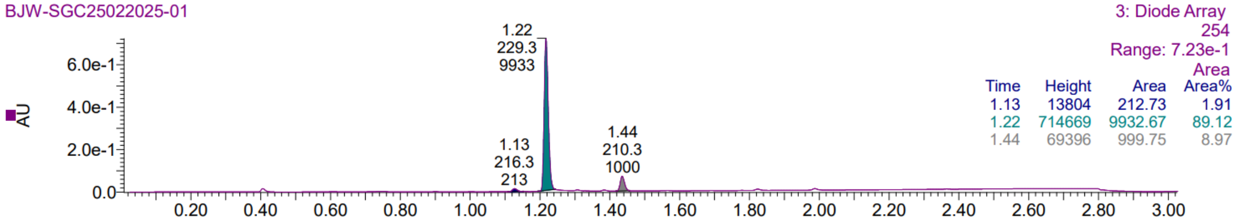


**2** - Batch B:

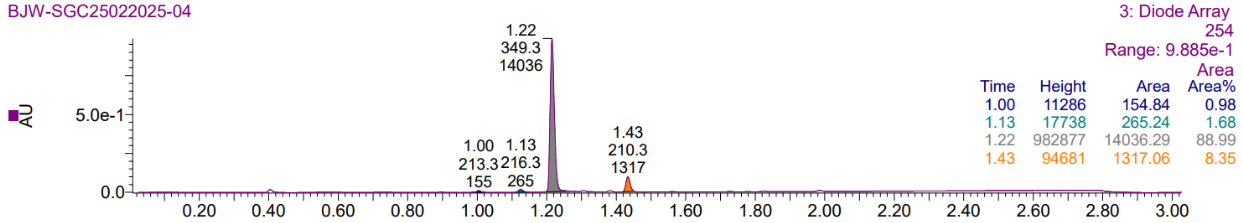

*2-(3,3-difluorocyclobutyl)-N-((1-methyl-1H-indazol-6-yl)methyl)aniline, **3***

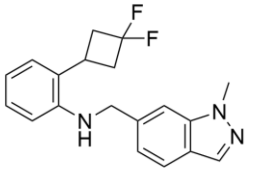


Chemical Formula: $C_{19}H_{19}F_2N_3$
Exact Mass: 327.15
Molecular Weight: 327.38

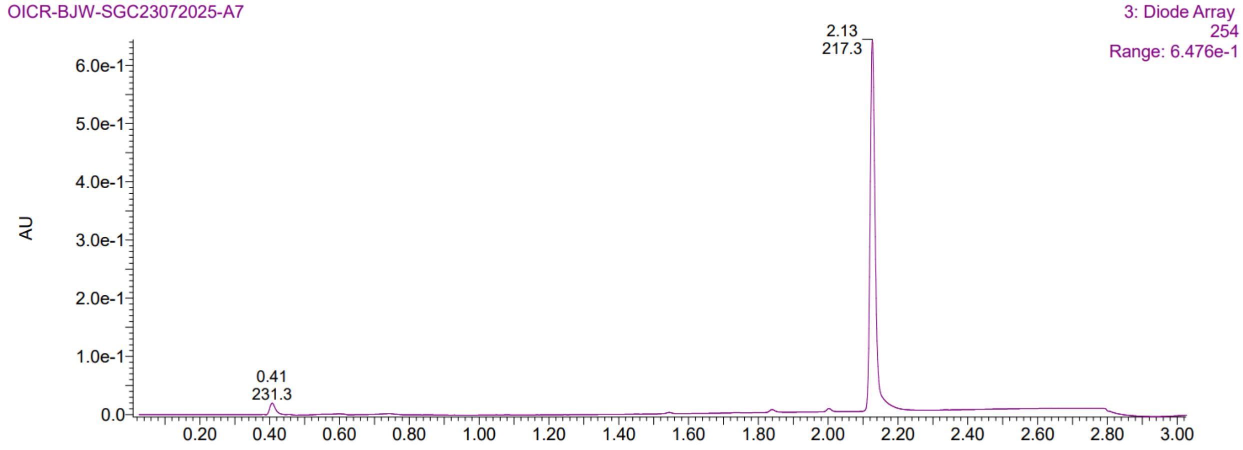


*2-cyclopropyl-N-((1-methyl-1H-indazol-6-yl)methyl)aniline, **4***

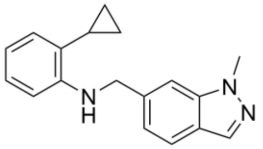


Chemical Formula: $C_{18}H_{19}N_3$
Exact Mass: 277.16
Molecular Weight: 277.37

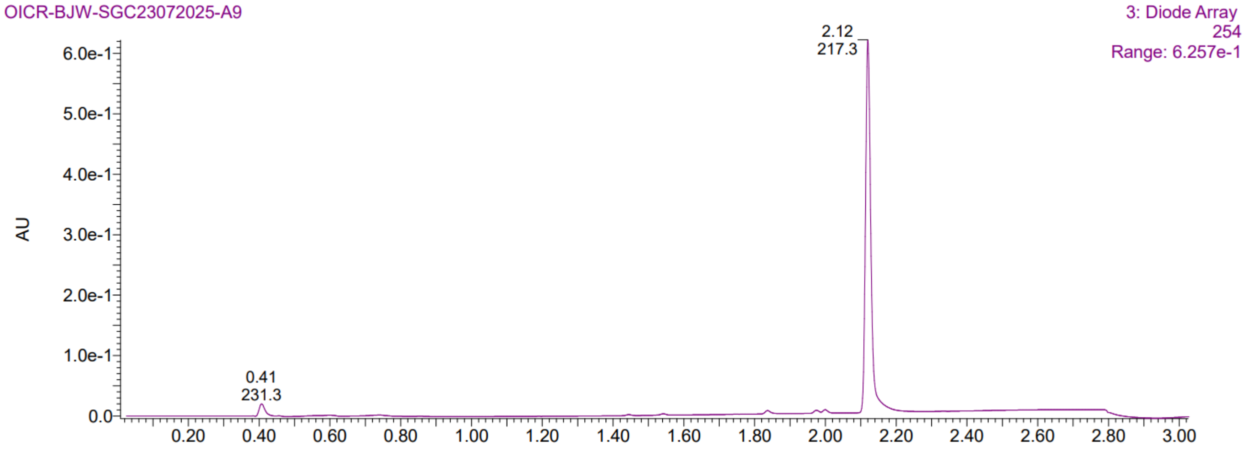

*2-(2,2-difluorocyclobutyl)-N-((1-methyl-1H-indazol-6-yl)methyl)aniline,* ***5***

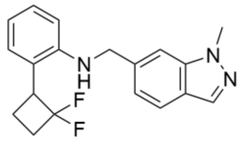


Chemical Formula: $C_{19}H_{19}F_2N_3$
Exact Mass: 327.15
Molecular Weight: 327.38

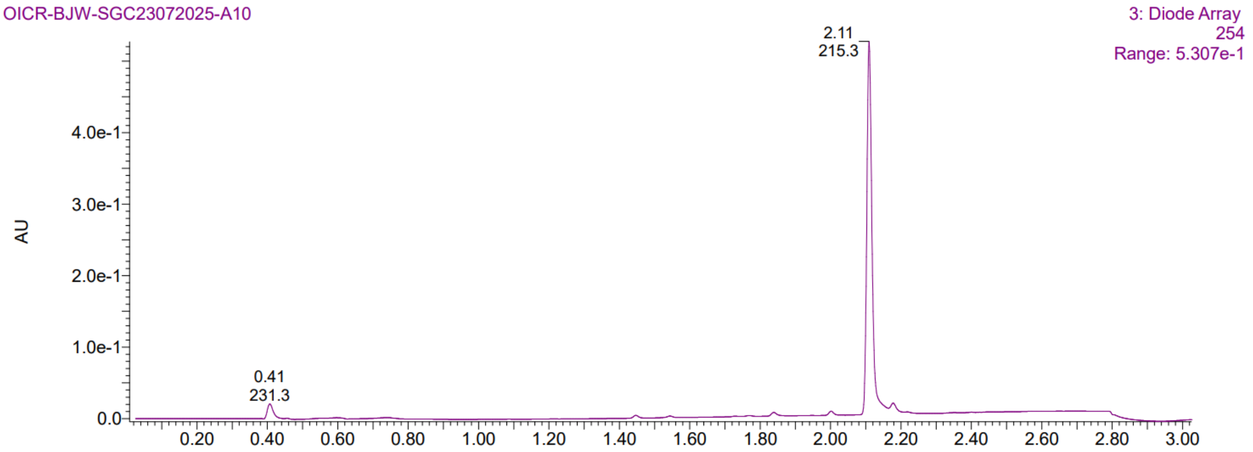


*N-((1-methyl-1H-indazol-6-yl)methyl)-2-(tetrahydrofuran-3-yl)aniline,* ***6***

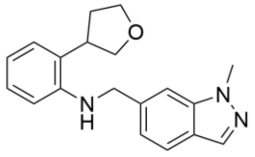


Chemical Formula: $C_{19}H_{21}N_3O$
Exact Mass: 307.17
Molecular Weight: 307.40

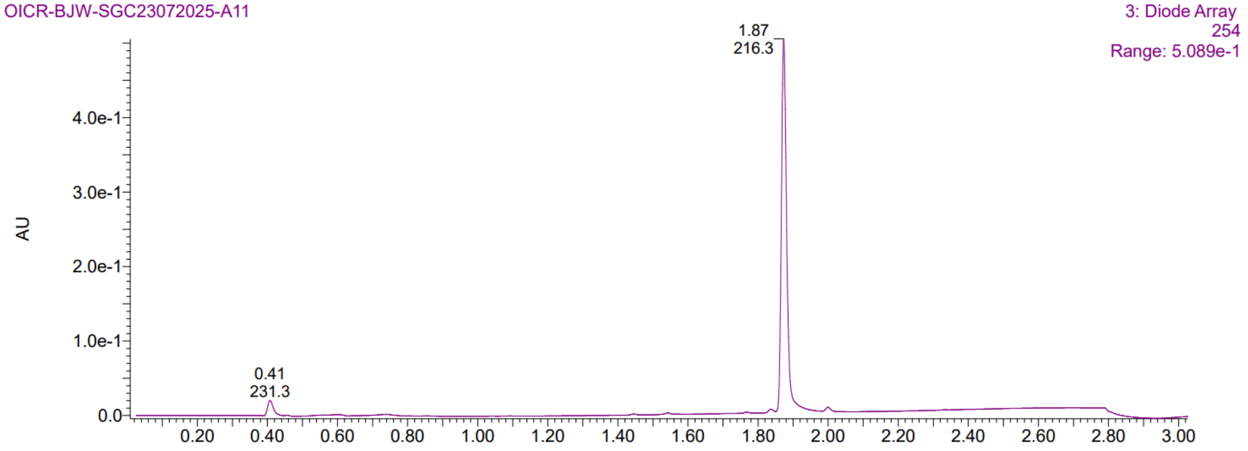

*N-((1-methyl-1H-indazol-6-yl)methyl)-2-(1,4-dioxaspiro[4.5]decan-8-yl)aniline, **7***

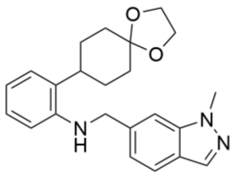

Chemical Formula: $C_{23}H_{27}N_3O_2$
Exact Mass: 377.21
Molecular Weight: 377.49

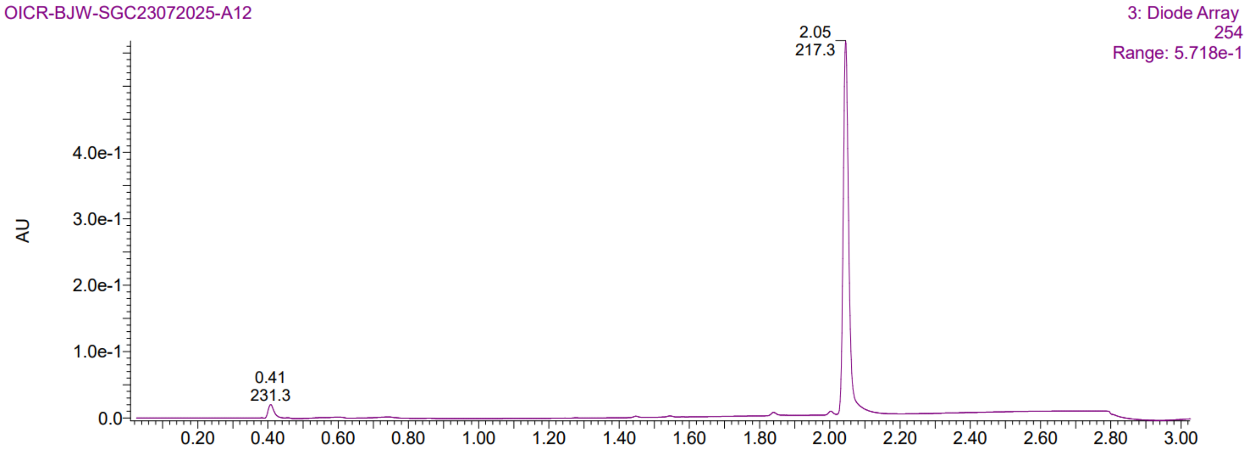


*tert-butyl 2-(2-(((1-methyl-1H-indazol-6-yl)methyl)amino)phenyl)acetate, **8***

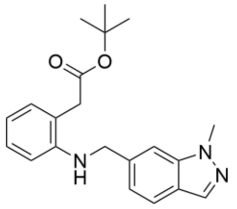

Chemical Formula: $C_{21}H_{25}N_3O_2$
Exact Mass: 351.19
Molecular Weight: 351.45

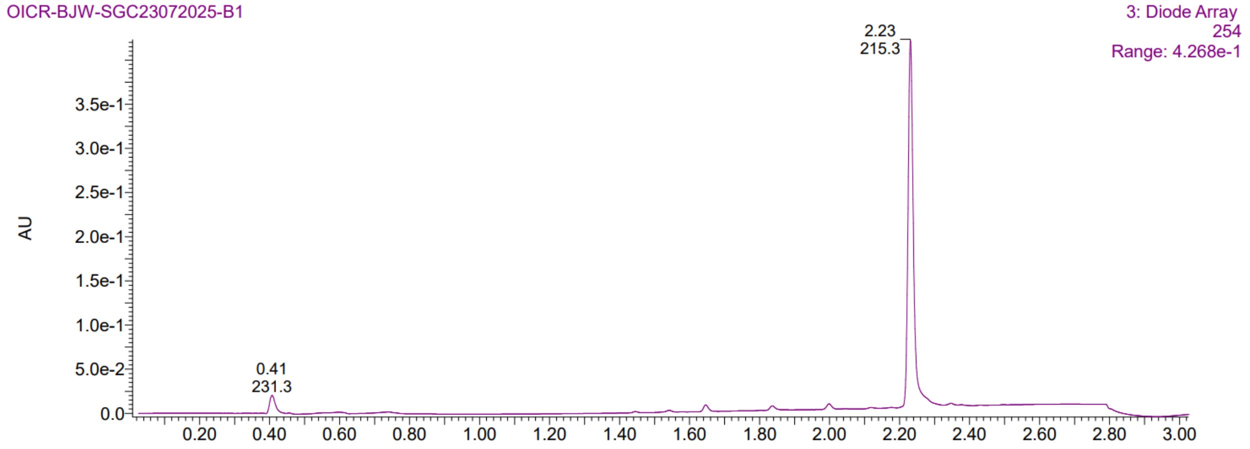

*2-(2,2-difluorocyclopropyl)-N-((1,3-dimethyl-1H-indazol-6-yl)methyl)aniline, **9***

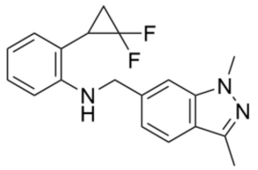


Chemical Formula: $C_{19}H_{19}F_2N_3$
Exact Mass: 327.15
Molecular Weight: 327.38

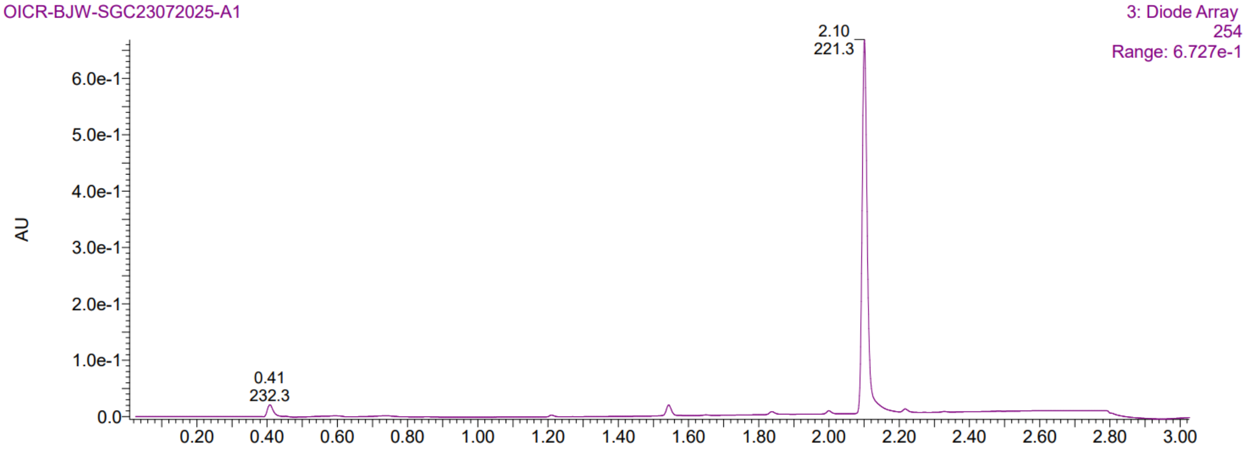


*2-(2,2-difluorocyclopropyl)-N-((1,4-dimethyl-1H-indazol-6-yl)methyl)aniline, **10***

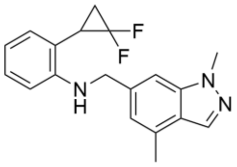


Chemical Formula: $C_{19}H_{19}F_2N_3$
Exact Mass: 327.15
Molecular Weight: 327.38

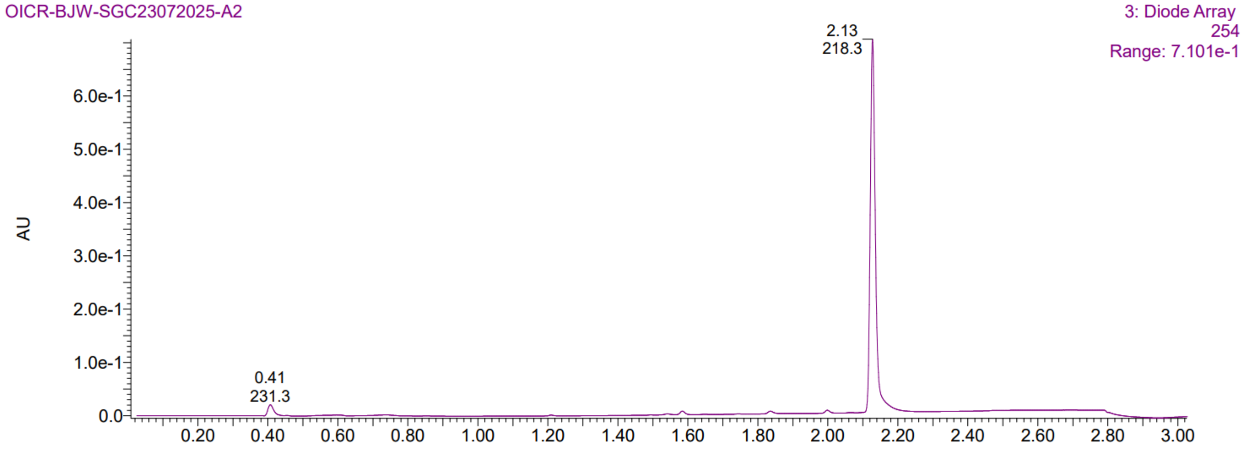

*2-(2,2-difluorocyclopropyl)-N-((1,5-dimethyl-1H-indazol-6-yl)methyl)aniline, **11***

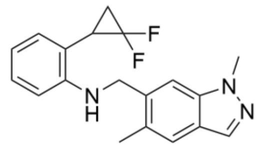


Chemical Formula: $C_{19}H_{19}F_2N_3$
Exact Mass: 327.15
Molecular Weight: 327.38

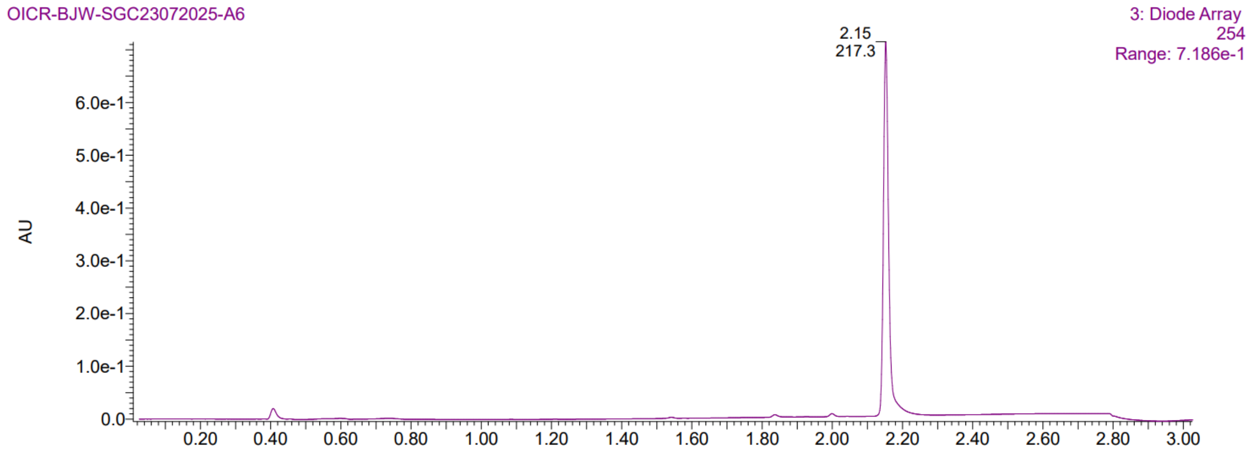


*2-(2,2-difluorocyclopropyl)-N-((2,4-dimethyl-2H-indazol-6-yl)methyl)aniline, **12***

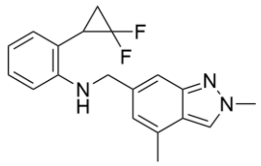


Chemical Formula: $C_{19}H_{19}F_2N_3$
Exact Mass: 327.15
Molecular Weight: 327.38

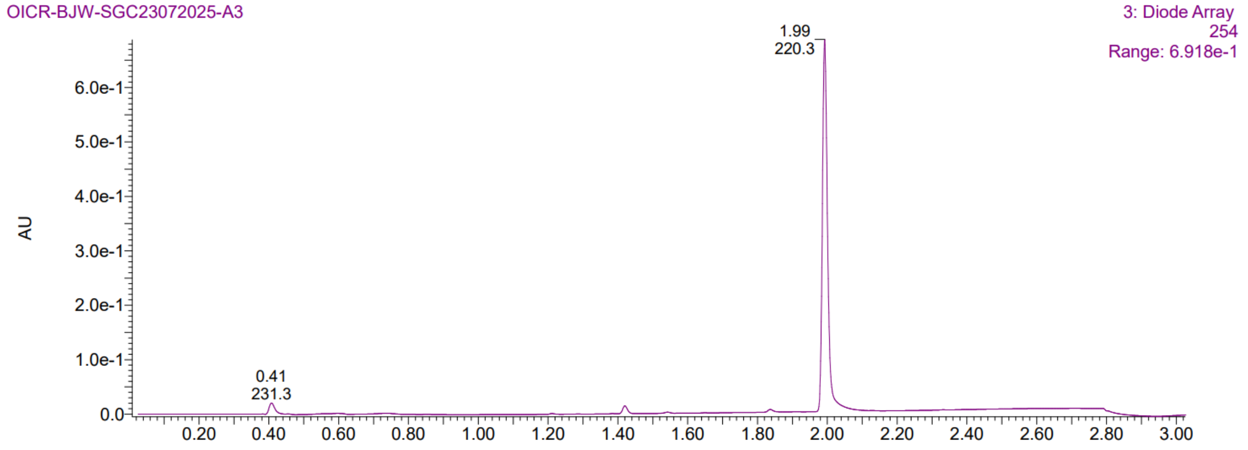

*2-(2,2-difluorocyclopropyl)-N-((2,3-dimethyl-2H-indazol-6-yl)methyl)aniline,* ***13***

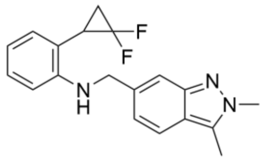


Chemical Formula: $C_{19}H_{19}F_2N_3$
Exact Mass: 327.15
Molecular Weight: 327.38

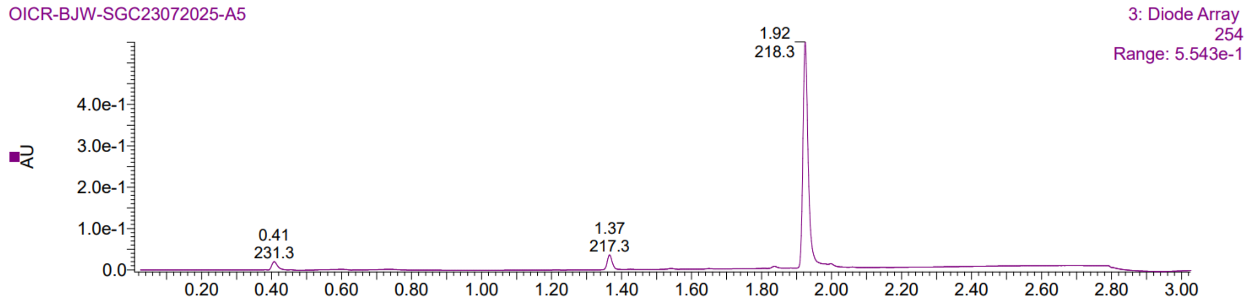


*2-(2,2-difluorocyclopropyl)-N-((2,5-dimethyl-2H-indazol-6-yl)methyl)aniline,* ***14***

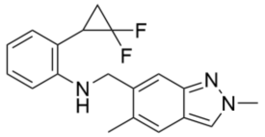


Chemical Formula: $C_{19}H_{19}F_2N_3$
Exact Mass: 327.15
Molecular Weight: 327.38

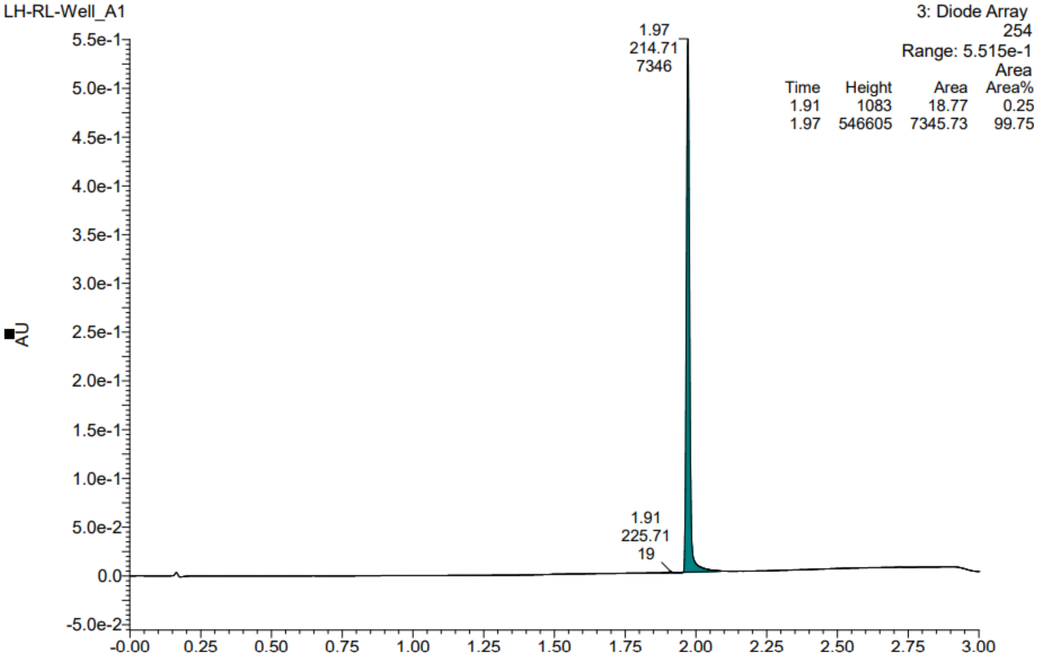

*2-(2,2-difluorocyclopropyl)-N-(1-(1-methyl-1H-indazol-6-yl)ethyl)aniline,* ***15***

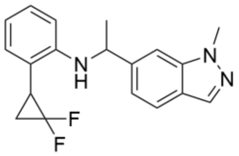

Chemical Formula: $C_{19}H_{19}F_2N_3$
Exact Mass: 327.15
Molecular Weight: 327.38

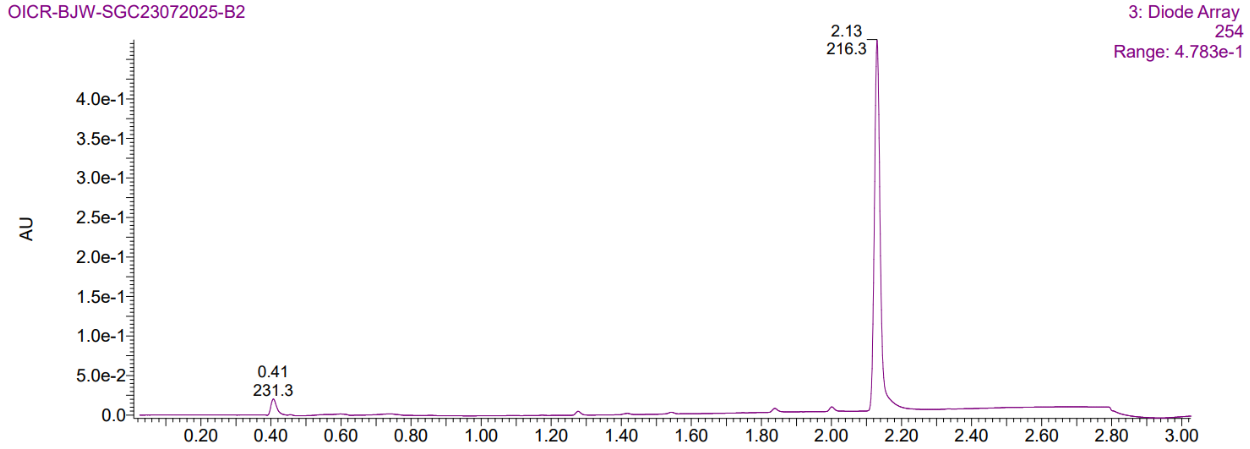


*N-(2-(2,2-difluorocyclopropyl)phenyl)-1-methyl-1H-indazole-6-carboxamide,* ***16***

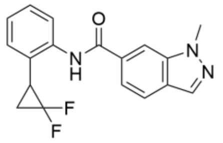

Chemical Formula: $C_{18}H_{15}F_2N_3O$
Exact Mass: 327.12
Molecular Weight: 327.33

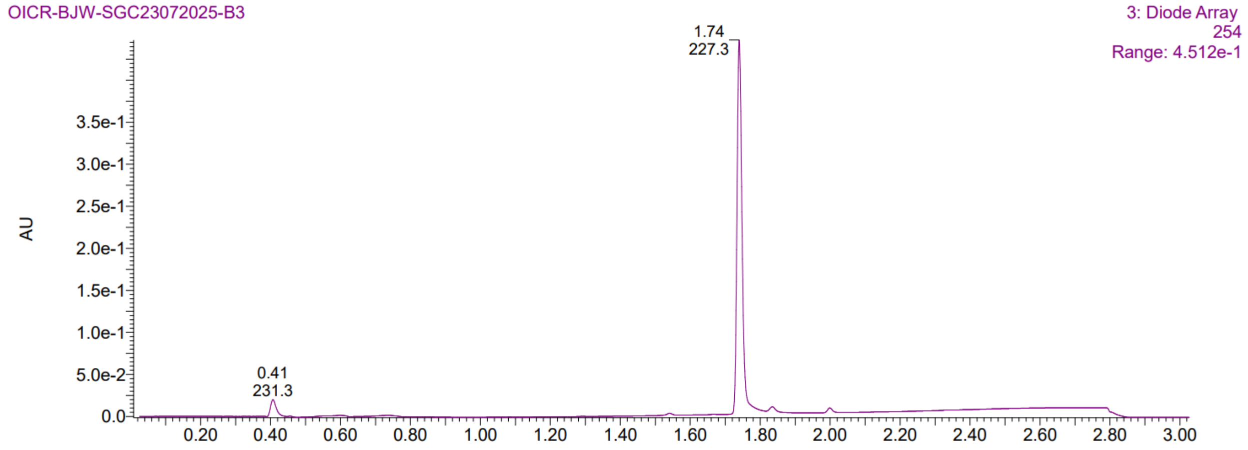

*2-(2,2-difluorocyclopropyl)-N-methyl-N-((1-methyl-1H-indazol-6-yl)methyl)aniline, **17***

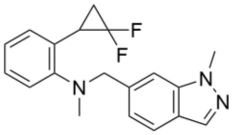

Chemical Formula: $C_{19}H_{19}F_2N_3$
Exact Mass: 327.15
Molecular Weight: 327.38

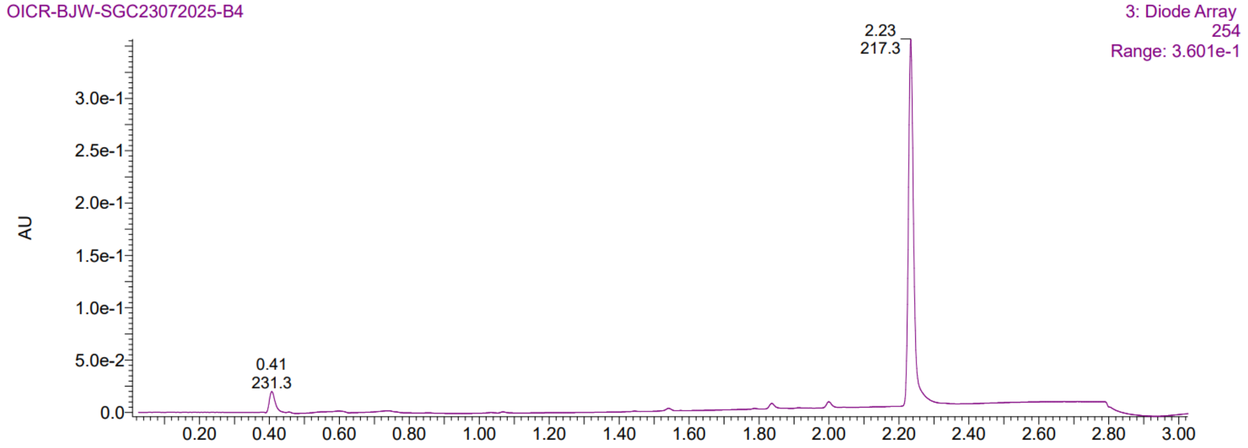


*2-(2,2-difluorocyclopropyl)-5-methyl-N-((1-methyl-1H-indazol-6-yl)methyl)aniline, **18***

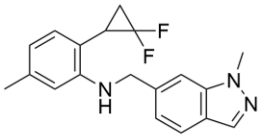

Chemical Formula: $C_{19}H_{19}F_2N_3$
Exact Mass: 327.15
Molecular Weight: 327.38

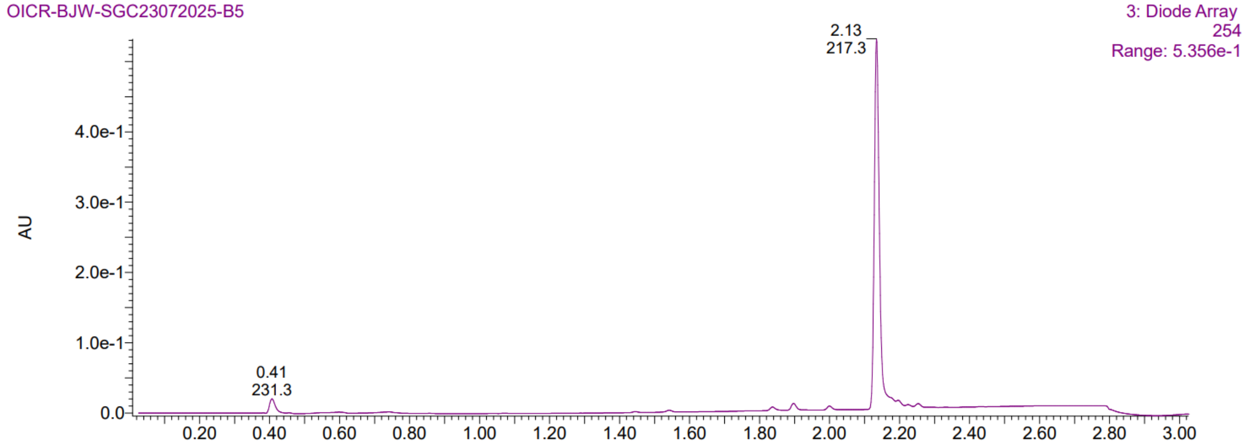

*3-(3,3-difluorocyclobutyl)-4-(((1-methyl-1H-indazol-6-yl)methyl)amino)phenol,* ***19***

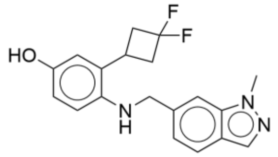

Chemical Formula: $C_{19}H_{19}F_2N_3O$
Exact Mass: 343.15
Molecular Weight: 343.38

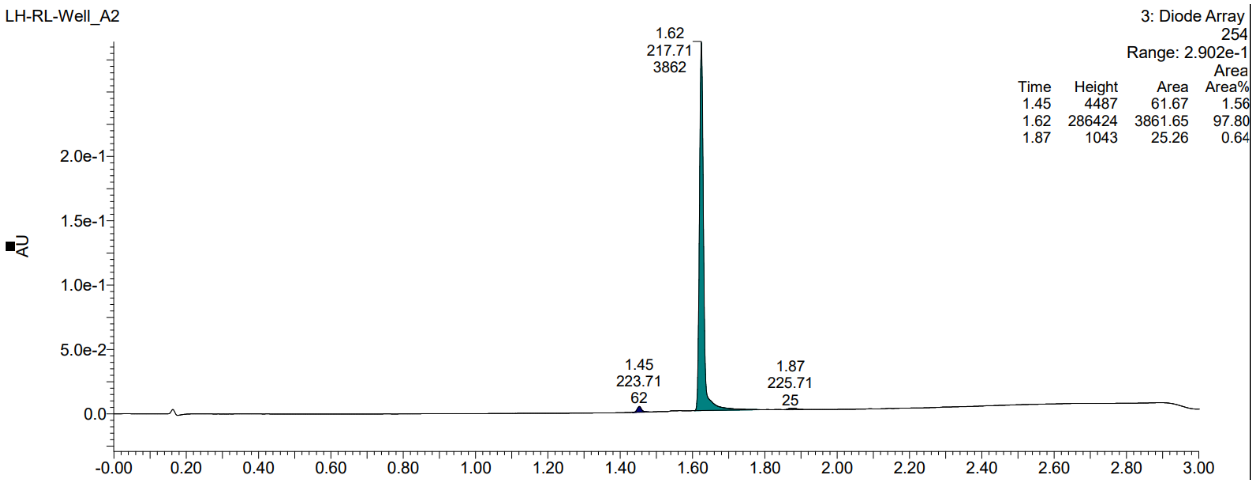


*2-(3,3-difluorocyclobutyl)-4-methoxy-N-((1-methyl-1H-indazol-6-yl)methyl)aniline,* ***20***

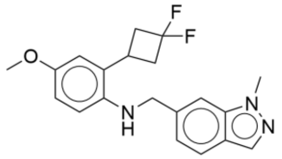

Chemical Formula: $C_{20}H_{21}F_2N_3O$
Exact Mass: 357.17
Molecular Weight: 357.40

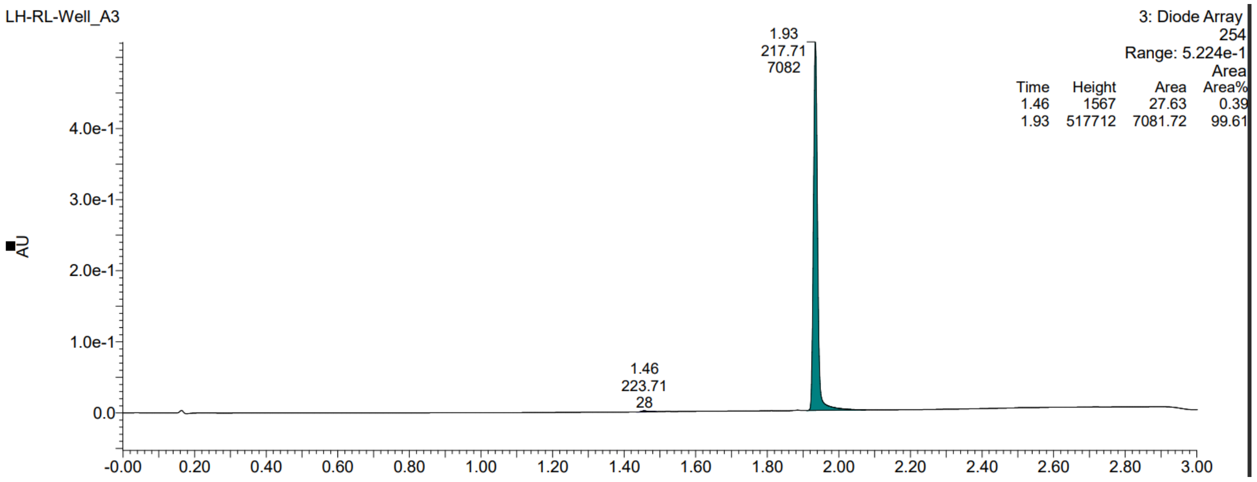

*2-(3,3-difluorocyclopentyl)-N-((1,3-dimethyl-1H-indazol-6-yl)methyl)aniline, **21***

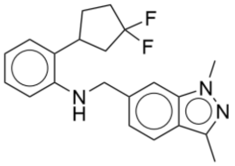

Chemical Formula: $C_{21}H_{23}F_2N_3$
Exact Mass: 355.19
Molecular Weight: 355.43

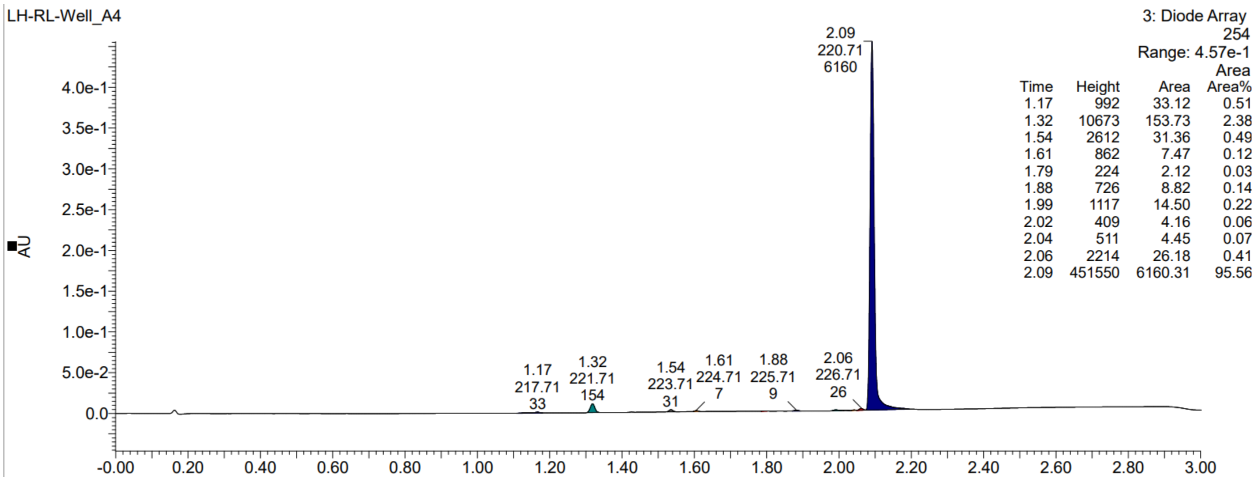


*2-(3,3-difluorocyclobutyl)-N-((3-fluoro-1-methyl-1H-indazol-6-yl)methyl)aniline, **22***

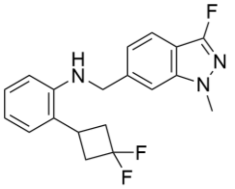

Chemical Formula: $C_{19}H_{18}F_3N_3$
Exact Mass: 345.15
Molecular Weight: 345.37

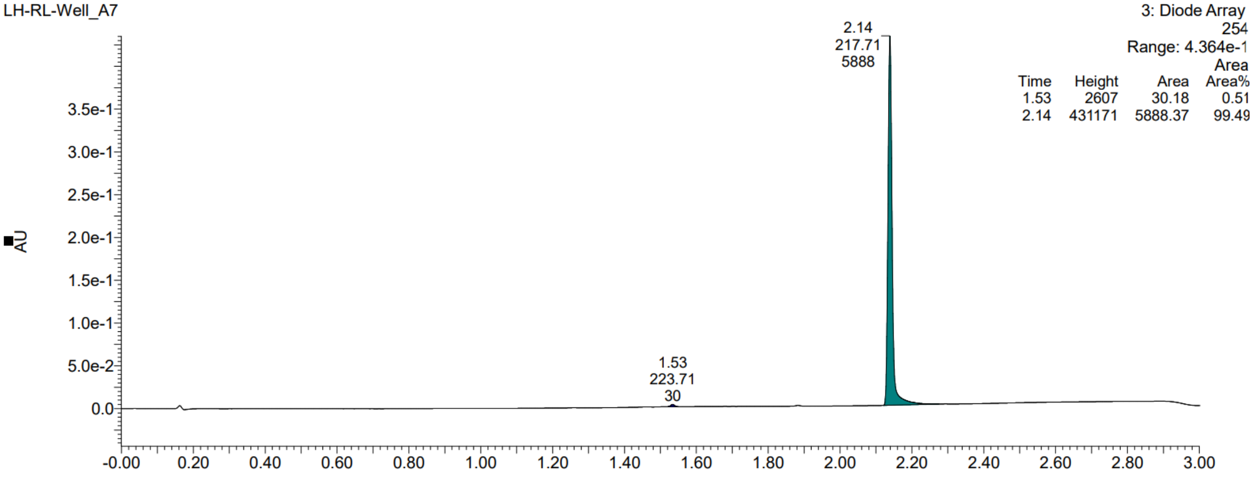

*N-((3-chloro-1-methyl-1H-indazol-6-yl)methyl)-2-(3,3-difluorocyclobutyl)aniline, **23***

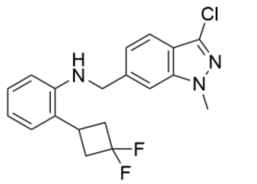

Chemical Formula: $C_{19}H_{18}ClF_2N_3$
Exact Mass: 361.12
Molecular Weight: 361.82

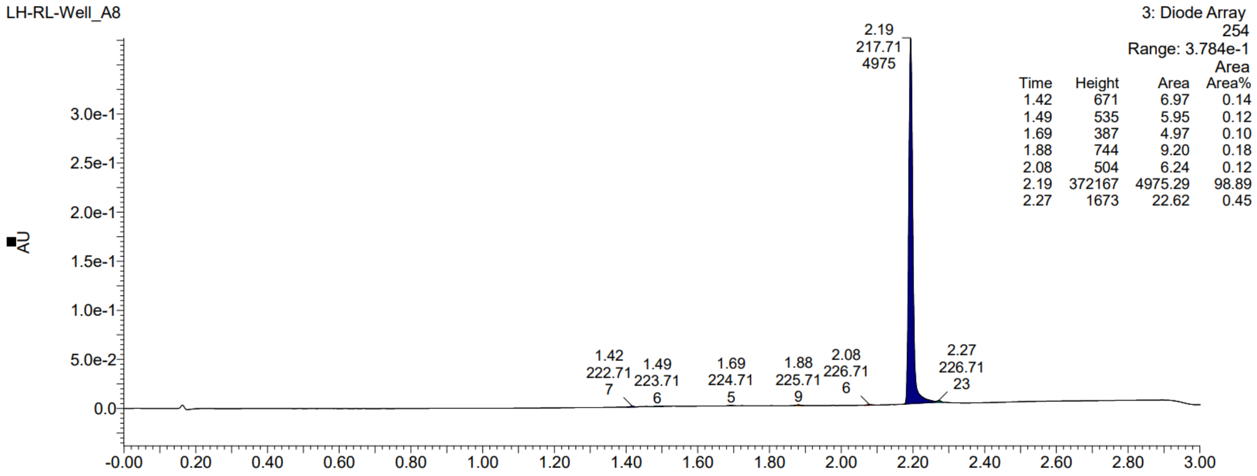


*N-((3-methyl-1H-indazol-6-yl)methyl)-2-(pyridin-2-yl)isoindolin-4-amine, **24***

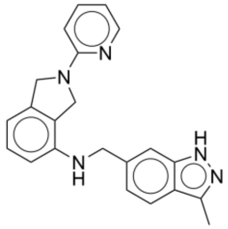

Chemical Formula: $C_{22}H_{21}N_5$
Exact Mass: 355.18
Molecular Weight: 355.44

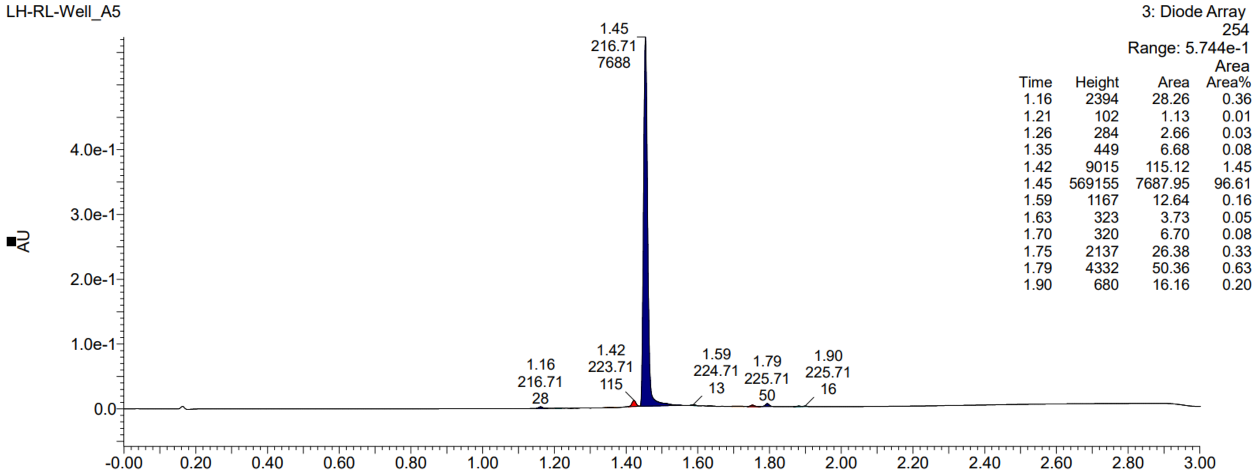

*3-(((3-amino-1H-indazol-6-yl)methyl)amino)-3'-(hydroxymethyl)-[1,1'-biphenyl]-4-carbonitrile,* ***25***

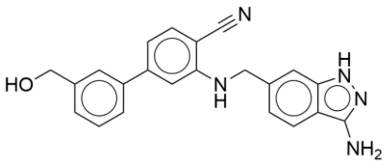


Chemical Formula: $C_{22}H_{19}N_5O$
Exact Mass: 369.16
Molecular Weight: 369.43

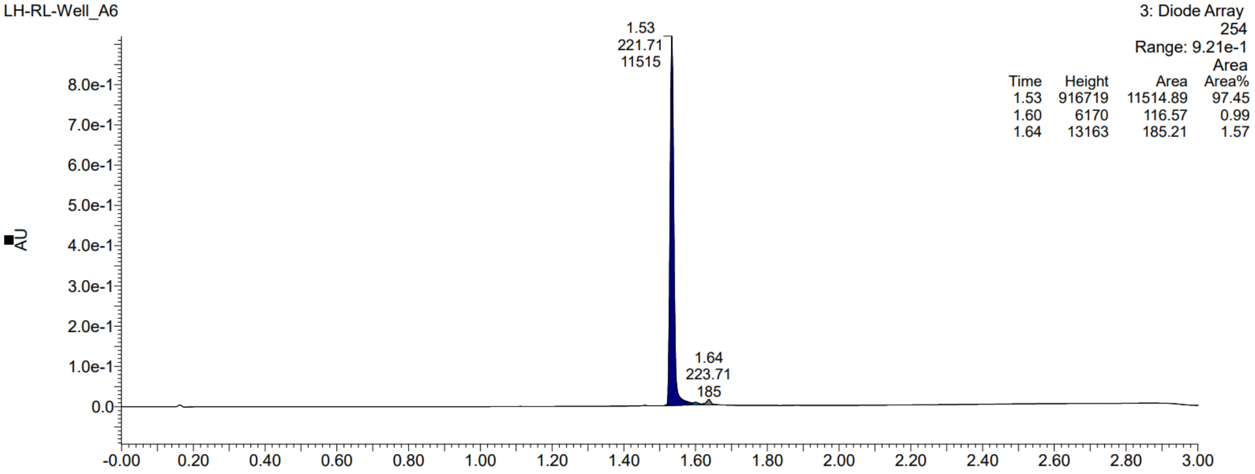
